# Host-induced gene silencing involves Arabidopsis ESCRT-III pathway for the transfer of dsRNA-derived siRNA

**DOI:** 10.1101/2020.02.12.945154

**Authors:** Schlemmer Timo, Weipert Lisa, Barth Patrick, Werner Bernhard Timo, Preußer Christian, Hardt Martin, Möbus Anna, Biedenkopf Dagmar, Claar Martina, Jelonek Lukas, Goesmann Alexander, Garikapati Vannuruswamy, Spengler Bernhard, Busche Tobias, Kalinowski Jörn, Koch Aline

**Affiliations:** Institute of Phytomedicine, University of Hohenheim, Otto-Sander-Strasse 5, D-70599, Stuttgart, Germany; Institute of Phytopathology, Centre for BioSystems, Land Use and Nutrition, Justus Liebig University, Heinrich-Buff-Ring 26, D-35392, Giessen, Germany; Institute of Bioinformatics and Systems Biology, Justus Liebig University, Heinrich-Buff-Ring 58, D-35392, Giessen, Germany; Center for Tumor Biology and Immunology, Philipps University, Hans-Meerwein Straße 3, D-35032, Marburg, Germany; Biomedical Research Centre Seltersberg (BFS), Imaging Unit, Justus Liebig University, Schubertstraße 81, D-35392, Giessen, Germany; Institute of Inorganic and Analytical Chemistry, Justus Liebig University, Heinrich-Buff-Ring 17, D-35392, Giessen, Germany; Center for Biotechnology - CeBiTec, Bielefeld University, Universitätsstraße 27, D-33615, Bielefeld, Germany

**Author notes:** shared first authorship (Schlemmer Timo, Weipert Lisa).

## Abstract

Small (s)RNAs and their double-stranded (ds)RNA precursors have been adopted to control diseases in crop plants through expression in transgenic plants and targeted gene silencing (host-induced gene silencing, HIGS). While HIGS strategies proved to be effective, the mechanism of RNA transfer at the plant - pathogen interface is widely unknown. Here we show that extracellular vesicles (EVs) purified from *Arabidopsis thaliana* plants expressing CYP3RNA, a dsRNA originally designed to target the three *CYP51* genes of the fungal pathogen *Fusarium graminearum*, contain CYP3RNA-derived small interfering (si)RNAs as shown by RNA sequencing (RNA-seq) analysis. These transgene specific siRNAs had a length of 21 and 22 nucleotides with a bias towards 5’-uracil (U) and 5’-adenine (A). Notably, stringent protease and RNase treated EV fractions contained >70% less CYP3RNA-derived siRNAs, suggesting the presence of co-purified extravesicular nucleoprotein complexes stabilizing siRNAs outside of EVs. In addition, mutants of the ESCRT-III complex showed a loss of HIGS-mediated disease resistance and EVs isolated from these mutants were free of CYP3RNA-derived siRNAs. Together, these findings support the view that endosomal vesicle trafficking is required for HIGS mediating the transfer of transgene-derived siRNAs between donor host cells and recipient fungal cells probably in an EV-independent manner.

## Introduction

Small (s)RNA effectors play a crucial role in the outcome of plant-pathogen interactions (Hua et al. 2018, Huang et al. 2019, Zeng et al. 2019) and have a great potential for controlling diseases and pests in crop plants (Cai et al. 2018a, Koch and Kogel, 2014, Wang et al. 2017a). A multitude of transgenic crops expressing double-stranded (ds)RNAs targeting essential genes through RNA interference (RNAi) are more resistant to viroids (Schwind et al 2009), viruses (Waterhouse et al. 1998), bacteria (Escobar et al. 2001)], fungi (Chen et al. 2016, Cheng et al. 2015, Koch et al. 2013, Panwar et al. 2018), oomycetes (Govindarajulu et al. 2015), nematodes (Chaudhary et al. 2019, Shivakumara et al. 2017), and insects (Coleman et al. 2015, Abdellatef et al. 2015). Host-induced gene silencing (HIGS) involves *i.* processing of transgene-derived dsRNA into small interfering (si)RNAs, *ii.* siRNA transfer into the interacting microbe/pest, and *iii.* siRNA interference with their target transcripts, though many details of the process have not been fully elucidated. Several reports suggested to use exogenous RNA to control fungal diseases (Cai et al. 2018b, Dalakouras et al. 2020, Dubrovina and Kiselev, 2019, Koch et al. 2016, Mitter et al, 2017). In the case of spray-induced gene silencing (SIGS) both the plant and the microbe silencing machinery appears to contribute to inhibition of the fungus (Koch et al. 2016).

Despite the promising prospective for RNAi-based disease control in agriculture, little is known about the mechanisms underlying the transport of RNAs from the plant host to an interacting microbial pathogen. While nematodes and some insects possess a transmembrane channel-mediated dsRNA uptake mechanism for intake and cell-to-cell distribution based on *Caenorhabditis elegans* SID proteins (SYSTEMIC RNA INTERFERENCE DEFICIENT) (Huvenne and Smagghe, 2010), fungi and plants lack these proteins. Alternatively, it has been hypothesized that plant sRNA effectors are associated with RNA-binding proteins (RBPs) and/or extracellular vesicles (EVs) (Cai et al. 2018a, Rutter and Innes; 2018). The interface between a plant and an interacting fungus is the primary site for mutual recognition, where vesicle secretion from both organisms occurs (An et al. 2006a, An et al. 2006b, Albuquerque et al. 2008). Several types of EVs, defined according to size and origin, have been identified in eukaryotic cells (Gould and Raposo, 2013). The largest class are apoptotic bodies with diameters of 800-5,000 nm. They are released by cells undergoing programmed cell death, thus reflecting the initially anticipated role of EVs as waste disposal system. EVs within a size range of 100-1,000 nm are categorized as large EVs that are released by budding from the plasma membrane (PM), while endosome-derived small EVs instead are only 30-200 nm in diameter and are released when multivesicular bodies (MVBs) fuse with the PM (Akers et al. 2013, Colombo et al, 2014, György et al. 2011, Liegeois et al. 2006, van der Pol et al 2012). In mammals, exosomes mediate intercellular communication by shuttling proteins, lipids and RNAs, where RNA molecules remain functional after delivery and can elicit effects in the recipient cell (Mittelbrunn et al. 2011, Pegtel et al. 2010, Ridder et al. 2014).

Based on this knowledge, we speculated that HIGS-mediating RNAs also are loaded into plant vesicles and transferred by EVs that cross the plant-fungus interface. It was reasoned already more than 50 years ago that a fusion of plant MVBs with the PM may result in the release of small vesicles into the extracellular space (Halperin and Jensen, 1967). Consistent with this early observation, biogenesis of immune-related MVBs (An et al. 2006a, An et al. 2006b) and the release of their cargo via EVs also is inherent to the plant secretory pathway (Rutter and Innes, 2017). Transmission electron microscopy (TEM) revealed proliferation of MVBs next to plant cell wall papillae in response to infection with powdery mildew fungus and the release of para-mural vesicles. The authors proposed that released vesicles might be similar to exosomes in animal cell thus anticipating that exosomes exist in plants (An et al. 2006a, An et al. 2006b). Immune-related plant EVs may carry defence compounds to strengthen the plant cell wall at the site of fungal attack. Supporting this notion, proteins, hydrogen peroxide, and callose could be identified inside MVBs next to the PM (An et al. 2006b, Xu and Mendgen 1994). EVs also were identified in the extrahaustorial matrix of powdery mildew fungus, though it could not be determined whether these vesicles were of plant or fungal origin (Micali et al. 2011).

Rutter and Innes (Rutter and Innes, 2017) isolated EVs from endosomal origin in a size range of 50-300 nm from the apoplast of Arabidopsis leaves, thus providing a direct proof that EVs also exist in plants. Vesicle secretion was enhanced in response to either the bacterial pathogen *Pseudomonas syringae* or the defence signalling compound salicylic acid. Consistent with this, the EV proteome was enriched for proteins associated with abiotic and biotic stress responses, including proteins involved in signal transmission, such as RPM1-INTERACTING 4 (RIN4), proteins of the myrosinase-glucosinolate system, such as the glucosinolate transporters PENETRATION 3 (PEN3) shown to accumulate around powdery mildew haustoria (Stein et al, 2006), proteins involved in reactive oxygen species (ROS) signaling and oxidative stress responses, proteins involved in membrane-trafficking, among them syntaxins such as PEN1, and proteins for the transport of ions, water, and sugar substrates. Moreover, EVs contained other trafficking proteins such as RAB GTPases and PATELLIN 1 and 2 (PATL1 and PATL2). PATL1 and PATL2 bind phosphoinositides and mediate vesicle transport and/or fusion (Petersen et al. 2014). PATL1 localizes to the cell plate in dividing cells but also associates with PLASMODESMATA-LOCATED PROTEIN1 (PDLP1) during downy mildew infection and may colocalize with PDLP1 at the extrahaustorial membrane (Caillaud et al, 2014).

Here, we assess whether artificial, antifungal dsRNA originating from an endogenous transgene are incorporated into plant EVs. We show that EVs from leaves of transgenic Arabidopsis plants expressing CYP3RNA contain siRNAs derived from CYP3RNA, a 791 nt long noncoding dsRNA that concomitantly silences the two fungal cytochrome P450 sterol 14α-demethylase genes *CYP51A* and *CYP51B* and the virulence factor *CYP51C* thereby inhibiting the growth of *Fusarium graminearum* (*Fg*) both *in vitro* and *in planta* (Koch et al, 2013, Koch et al 2016, Koch et al, 2018). Moreover, mutants of the secretory vesicle pathway are strongly compromised in HIGS-mediated resistance and EVs isolated from these mutants were free of CYP3RNA-derived siRNAs.

## Results

### Exosome-like vesicles from leaf extracts of CYP3RNA-expressing transgenic Arabidopsis contain CYP3RNA-derived siRNAs

The current knowledge on plant EVs is consistent with the idea of an EV-mediated effector shuttle (Cai et al. 2018b, Dubey et al. 2019, Dunker et al. 2020, Wang et al. 2017b, Weiberg et al. 2013, Zanini et al. 2021, Zhang et al. 2012, Zhang et al. 2016, Zhu et al. 2017). Here, we tested the possibility that HIGS also involves vesicle transport of dsRNA-derived sRNAs. To this end, we isolated vesicles from plants expressing the 791 nt CYP3RNA that was previously shown to confer resistance against *Fg* by silencing the three fungal genes *FgCYP51A, FgCYP51B* and *FgCYP51C* (Koch et al. 2013, Koch et al. 2016, Koch et al, 2018). After differential centrifugation of the leaf cell extract (see MM), vesicles were pelleted by 160,000 xg ultracentrifugation and subsequently loaded on top of a sucrose density gradient. After an additional ultracentrifugation step, two green coloured bands in the gradients of samples from wt and CYP3RNA-expressing plants contained vesicles (Figure 1A, 1B). The upper band (fraction F160-s1) located between the 30% and 45% sucrose layer, and the lower band (fraction F160-s2), located between the 45% and 60% layer, were harvested separately and inspected by negative staining and TEM. Fractions F160-s1 contained vesicles that were in a size range of about 50 nm to 200 nm. Most of them were cup-shaped and surrounded by a layer reminiscent of a membrane (Figure 1A, 1B). Fractions F160-s2 contained only a few vesicles and most of them were not surrounded by a membrane (Supplementary Figure 1A), while the fraction between the two visible greenish bands was mainly free of vesicles (Supplementary Figure 1B). The size of the vesicles from the upper fraction was further determined using ImageJ, revealing an average diameter of about 95 nm for wt plants and 105 nm for CYP3RNA-expressing plants (Table 1). This size fits with the size definition of mammalian small EVs having 30 nm to maximal 150 nm in diameter and Arabidopsis EVs having 50 nm to maximal 300 nm in diameter (Colombo et al. 2014, Rutter and Innes 2017).

**Figure 1.**
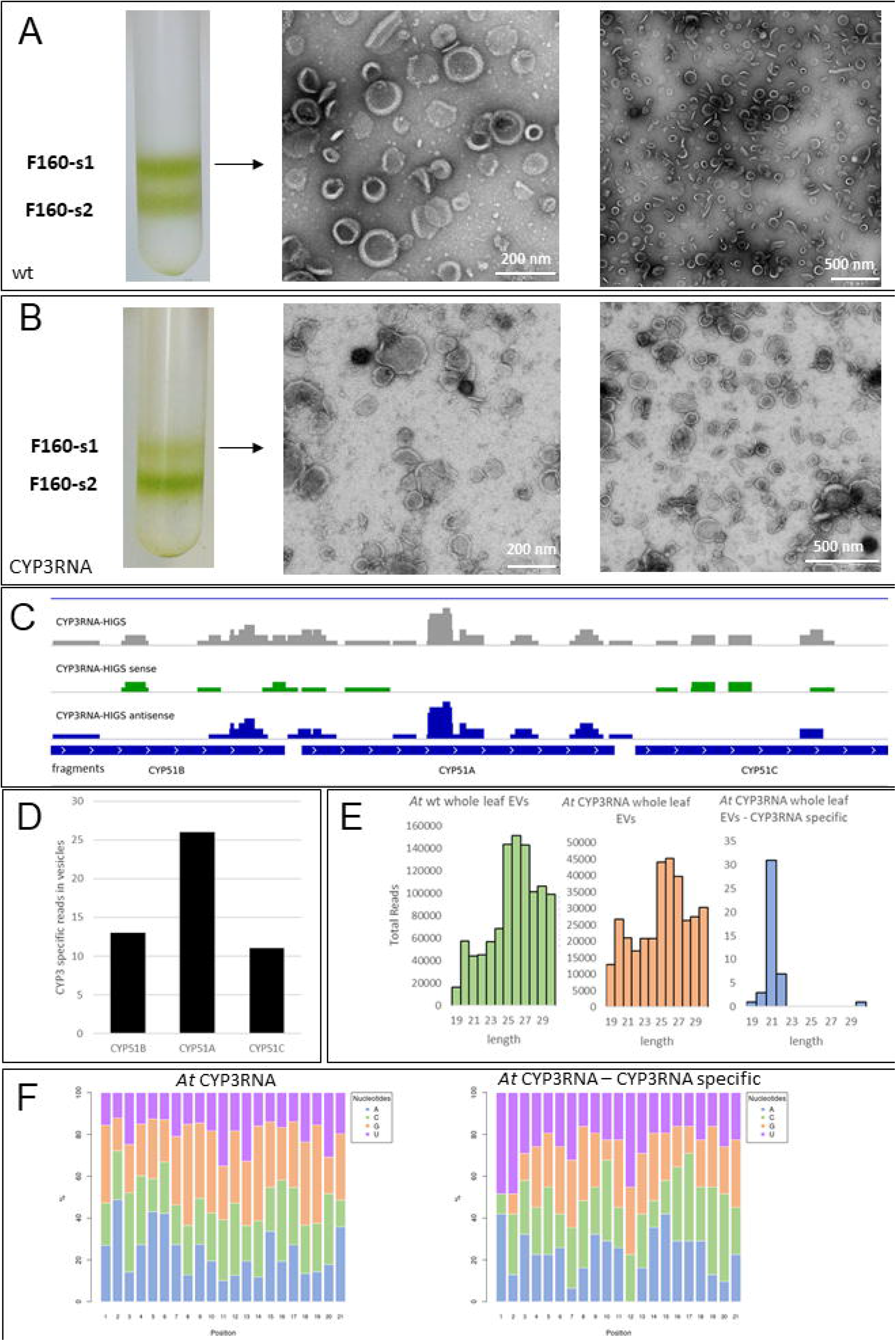
Vesicle isolation from Arabidopsis whole leaf cell extracts purified by sucrose-gradient centrifugation. (A)(B) Vesicles from wt Col-0 (A) and CYP3RNA-expressing (B) plants were subjected to sucrose gradient centrifugation at 160,000 *g* resulting in two predominant greenish bands between the 30% - 45% (upper band, F160-s1) and 45% - 60% (lower band, F160-s2). Vesicles were analysed by negative staining and TEM. The fraction F160-s1 contained vesicles with a size range of 50-200 nm. Most of them were cup-shaped and surrounded by a layer reminiscent of a membrane. (C) Profiling of CYP3RNA-derived siRNAs by sRNA-seq. Total RNAs were isolated from vesicles of fraction F160-s1 containing vesicles with average sizes of 95 nm and 105 nm (see Table 1). sRNA reads of max. 25 nt from CYP3RNA-expressing (CYP3RNA-HIGS) and wt plants were mapped to CYP3RNA that represents sequences from gene fragments of the *Fusarium graminearum* genes *FgCYP51A, FgCYP51B* and *FgCYP51C* (Koch et al, 2013, Koch et al, 2016). Sequencing data are gained from two pooled separate vesicle isolations experiments. (D) sRNA profiling of vesicles isolated from whole leaf cell extracts of CYP3RNA-expressing Arabidopsis plants. There is a bias of siRNA reads mapping to the CYP3RNA sequence as most reads map to the *FgCYP51A* fragment of the precursor. (E) Size distribution of total sRNA-reads isolated from wt plants, CYP3RNA plants and reads matching to the CYP3RNA precursor. (F) Nucleotide distribution of unique 21 nt sRNAs isolated from apoplastic washes of wt plants, CYP3RNA plants, and reads of CYP3RNA plants with perfect complementarity to the CYP3RNA precursor.

**Table 1.** Average diameter of vesicles from Arabidopsis. Average diameter was analysed with the ImageJ Software based on microscopy scale bars. Standard errors (SE) are given in parentheses. Arabidopsis CYP3, CYP3RNA-expressing plants.

| Plant | Vesicle diameter [nm] | Source |
| --- | --- | --- |
| Arabidopsis wt | 94.8 (± 2.5) | whole leaves |
| Arabidopsis CYP3 | 105.2 (± 3.6) | whole leaves |
| Arabidopsis wt | 118.6 (± 6.5) | apoplastic washes |
| Arabidopsis CYP3 | 141.4 (± 8.0) | apoplastic washes |

Next, we assessed whether vesicles from leaf extracts of transgenic Arabidopsis contain RNA derived from the CYP3RNA precursor/transgene. Vesicles from the F160-s1 fraction were washed in 1x phosphate buffer saline (PBS), pelleted by 160,000 x*g* and RNA extraction was performed subsequently, with a final profiling of the sRNA by RNA-seq (sRNA-seq). Vesicles of the F160-s1 fractions yielded 720 ng (CYP3RNA plants) and 1,080 ng (wt plants) RNA, respectively, (purified each from 120 leaves (Supplementary Table 1)). Indexed sRNA libraries were constructed, pooled and sequenced on an Illumina MiSeq Platform, resulting in 5.3 and 2.1 million reads from libraries of CYP3RNA-expressing and wt plants, respectively (Supplementary Table 2). By mapping reads to the CYP3RNA precursor sequence, we found 31 unique siRNAs aligning to CYP3RNA (Figure 1C), while no CYP3RNA matching reads were found in vesicles from wt plants (Supplementary Figure 2). Overall, only 79,223 reads out of 5.3 million total reads from CYP3RNA plants were small RNAs (21 to 24 nts in length), which can be explained by the fact that total vesicle RNA was not size fractionated before sequencing. From the 31 siRNAs (total reads: 442 rpm) mapping to the CYP3RNA precursor sequence, 12 siRNAs (total reads: 177 rpm) matched the CYP51A fragment, 10 siRNAs (total reads: 126 rpm) matched the CYP51B fragment, and five siRNAs (88 rpm) matched the CYP51C fragment. Interesting to note, slightly more siRNAs matched to the CYP51A fragment which is in the middle of the CYP3RNA sequence (Figure 1D). EVs isolated from whole leaves contained most abundant sRNAs with a length of 25-27 nts for wt and CYP3RNA plants, while CYP3RNA transgene specific siRNAs were predominantly 21 and 22 nts in length (Fig. 1E). CYP3RNA-derived siRNAs had a bias towards 5’-uracil (U) and 5’-adenine (A), while the total sRNAs from wt plants and CYP3RNA plants showed no obvious bias towards a specific 5’nuleotide (Fig. 1F).

### EVs from apoplastic fluids of transgenic Arabidopsis contain CYP3RNA-derived siRNA

The above findings suggest that siRNAs from a transgene-derived dsRNA might be incorporated into plant vesicles. To further substantiate this finding, we isolated EVs from apoplastic washing fluids of CYP3RNA-expressing and wt Arabidopsis plants, following a published protocol (Rutter and Innes, 2017). Whole rosettes of 5-wk-old Arabidopsis plants were vacuum infiltrated with vesicle isolation buffer (VIB) until infiltration sites were visible as dark green leaf areas. After gathering apoplastic fluid by low-speed centrifugation, vesicles were isolated by differential centrifugation at 48,000 x*g* (F48) and 100,000 x*g* (F100). Pellets from the centrifugation steps were analysed by TEM. Consistent with the earlier report (Rutter and Innes, 2017), the F48 fraction contained vesicle-like structures resembling EVs in size and shape (Figure 2A). While the shape and structure of these vesicles were otherwise indistinguishable from vesicles isolated by sucrose gradient. Analysis of the particle diameter using ImageJ revealed an average diameter of about 141 nm and 119 nm for CYP3RNA-expressing and wt plants, respectively (Table 1). This fits well with the observation of Rutter and Innes (Rutter and Innes, 2017) showing that the most abundant particles in the F48 fraction had around 150 nm in diameter as detected by light-scattering. Also consistent with the above report, the F100 fraction was mostly devoid of vesicles and instead contained a multitude of small particles with diameters under 30 nm (Figure 2B). We further analysed the size of EVs by nanoparticle tracking analysis (NTA). We found a particle average size of 139 +/− 7.7 nm (Figure 2C). To further substantiate that the F40 fraction contains EVs, we conducted an immunoblot analysis detecting the membrane marker PEN1 (Rutter and Innes, 2017). We found a strong signal for PEN1 in extracts from the F40 fractions (Figure 2D).

**Figure 2.**
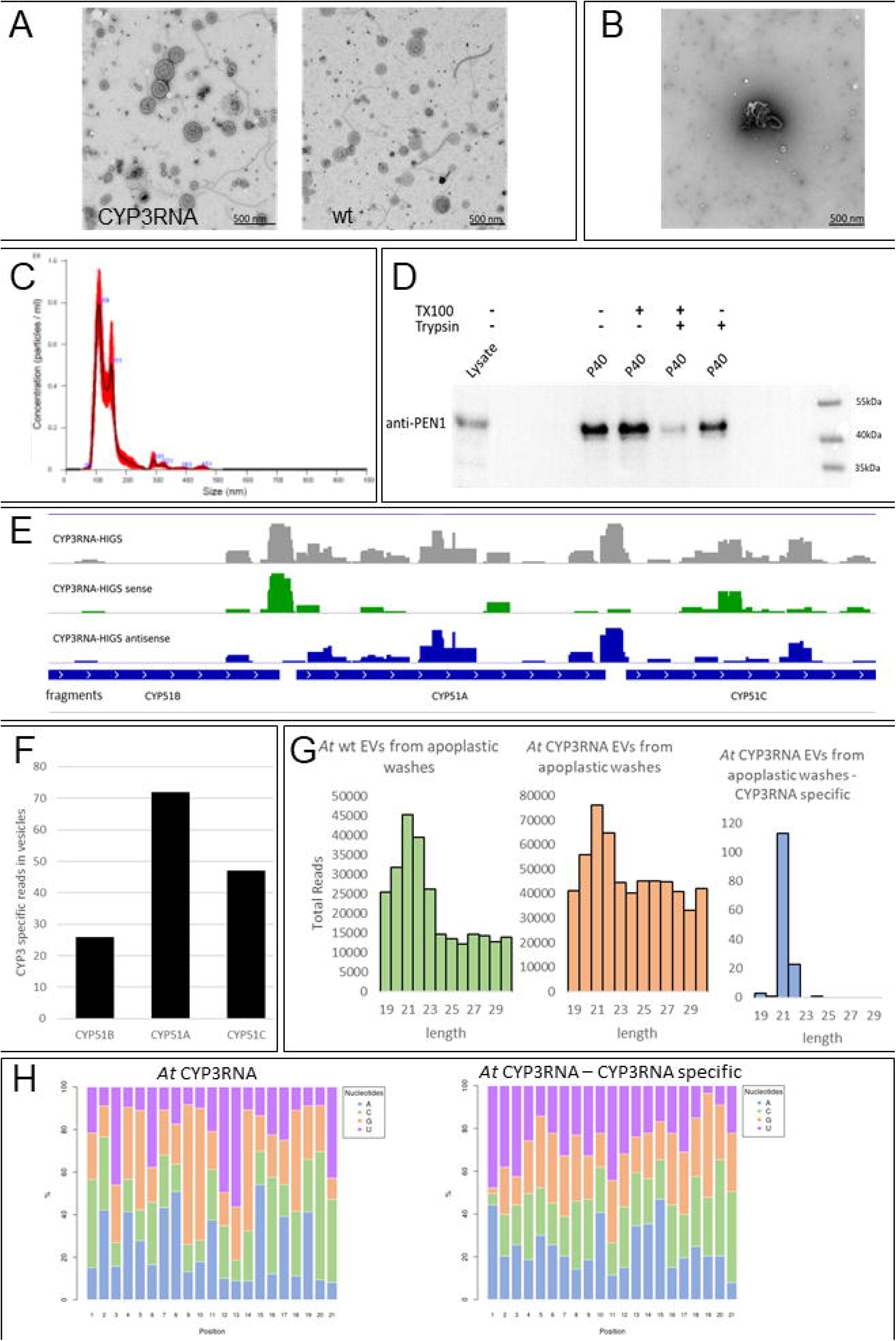
Analysis of EVs from the apoplastic fluid of Arabidopsis leaves. (A)(B) Negative staining and TEM of pellets of the 48,000 *g* (fraction F48, (A)) and the 100,000 *g* (F100, (B)) centrifugation of extracellular washing fluids. Pictures are shown exemplary for CYP3RNA-expressing and wt plants (A). Experiments were performed multiple times with similar results. (C) Size measurements by nanoparticle trafficking analysis (NTA) Averaged particle concentration per particle size purified from apoplastic washes of Arabidopsis CYP3RNA plants (black line) and +/− 1 standard error (red). Purified EVs have an average size of 139 +/7.7 nm the sample concentration was 1.0 x 10^10^ +/− 5.2 x 10^8^ particles/ml. (D) PEN1 detection was performed by immunoblot analysis. Proteins of whole cell extract (lysate) and isolated EVs were separated by electrophoresis and blotted on to a nitrocellulose membrane. Before separating, EVs were treated by Triton100, trypsin or both beside an untreated control. PEN1 was detected in all samples, but signal intensity was strongly reduced after EVs were treated by Triton100 and trypsin, while treatment by Triton100 or trypsin alone did not affect signal intensity. (E) Profiling of CYP3RNA-derived siRNAs in EVs from apoplastic fluid of Arabidopsis leaves. Vesicles were isolated by apoplastic washes. Total RNA extraction was performed from purified vesicles and analysed by sRNA-seq. sRNA reads of max. 25 nt from CYP3RNA-expressing (CYP3RNA-HIGS) plants were mapped to the sequence of CYP3RNA (Koch et al, 2013). Sequencing data are gained from vesicles of each 10 CYP3RNA-expressing plants. (F) Fragment distribution of CYP3RNA specific reads. (G) Read length distribution of sRNAs isolated from apoplastic washes of wt, CYP3RNA plants, and reads of CYP3RNA plants with perfect complementarity to the CYP3RNA precursor. (H) Nucleotide distribution of unique 21 nt sRNAs isolated from apoplastic washes of wt plants, CYP3RNA plants, and reads of CYP3RNA plants with perfect complementarity to the CYP3RNA precursor.

Next, we assessed the RNA cargo of wt and CYP3RNA plant derived EVs by RNAseq. From the total of 58 siRNA (607 rpm) mapping to CYP3RNA, 24 unique siRNAs (total reads: 242 rpm) mapped to the CYP51A fragment, four unique siRNAs (total reads: 31 rpm) mapped to the CYP51B fragment, and 19 siRNAs (total reads: 180 rpm) mapped to the CYP51C fragment, substantiating the slight bias for siRNAs matching the CYP51A fragment (Figure 2E, 2F). Notably, all EV samples revealed an accumulation of 21 and 22 nts siRNAs. Moreover, again we found CYP3RNA-derived siRNAs showing a bias towards a 5’-uracil (U) and 5’-adenine (A), while the total sRNAs from wt plants and CYP3RNA plants showed no obvious bias towards a specific 5’nuleotide (Figure 2G).

### Digestive treatments of EVs reveal high amounts of extravesicular RNAs and differ in their RNA composition

Isolation of plant EVs from apoplastic washes commonly results in copurification of extravesicular contaminants which can influence the interpretation of data (Rutter and Innes 2020). To overcome this limitation and to distinguish between siRNA from outside or inside EVs, we performed several EV treatments: protease and RNase to digest extracellular RNAs, proteins as well as ribonucleoprotein complexes. In addition, we treated EVs with Triton X (prior to RNase and protease digest) to break open EVs. From infiltration of at all 90 plants per genotype we gained 133 ng / 48 ng RNA (CYP3RNA/wt plants) from untreated EVs, 47 ng /47 (CYP3RNA/wt plants) ng from protease and RNase treated EVs and 24 ng / 42 ng (CYP3RNA/wt plants) from Triton X, protease and RNase treated EVs (Supplementary Table 1). Subsequently, RNA was subjected to sRNA-seq. Interestingly, we found that the RNA concentration and total reads decreases with each treatment step indicating 55-75% extravesicular RNAs for wt EVs and 70-85% for CYP3RNA plants compared to non-treated EV isolates (Figure 3A, 3B).

**Figure 3.**
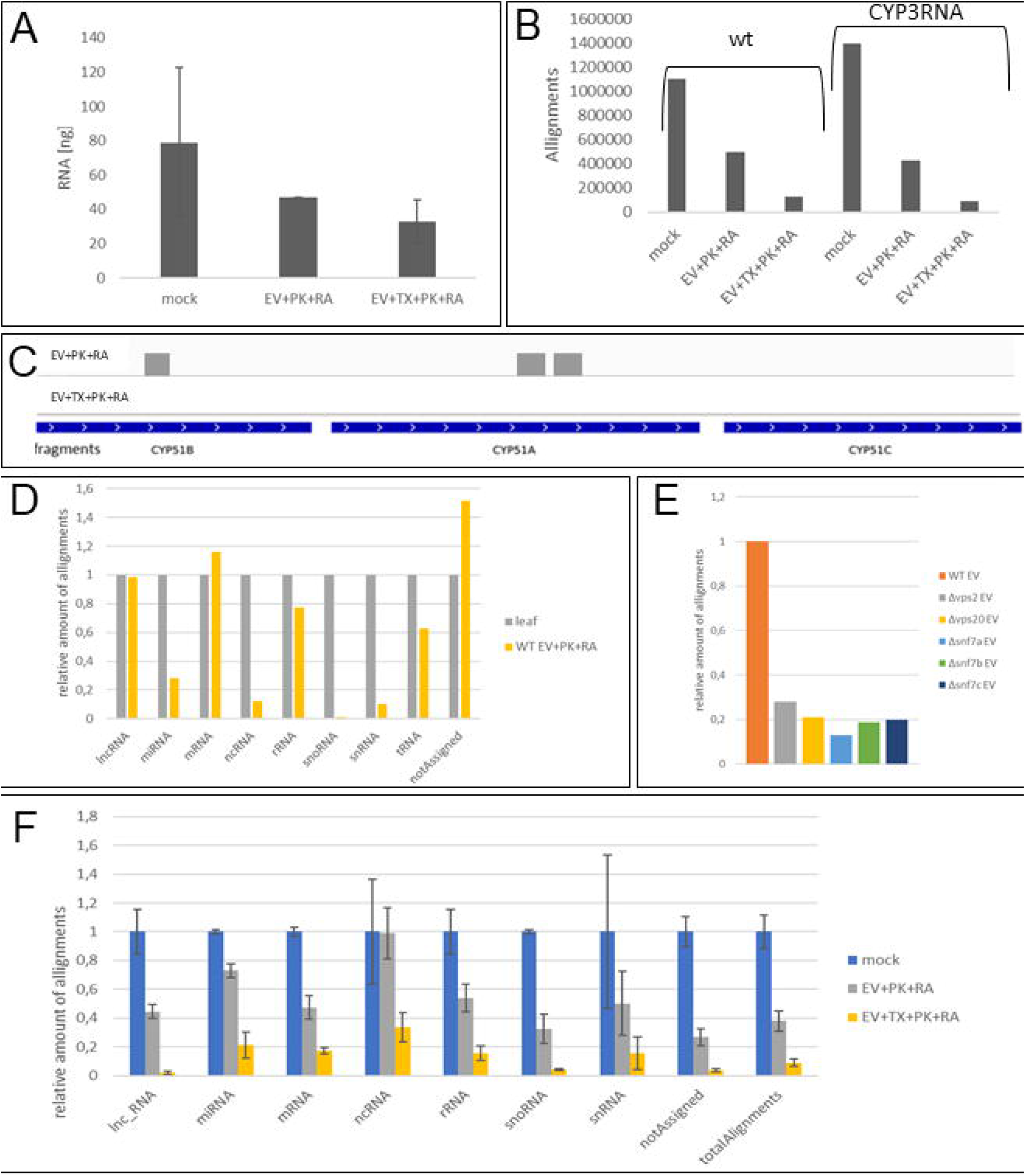
Analysis of the RNA composition of plant EVs. (A) Total amount of purified RNA from EVs of wt and CYP3RNA plants after differential treatments of the EVs. EVs were treated for 30 min at 37°C with protease K (PK), RNaseA (RA) and Triton-X 100 (TX) for EV disruption and cargo and contaminants digestion or PK and RA for contaminants digestion or only heat-treated under same conditions, which serves as a mock control (mock). (B) RNA from differential treated EVs from wt and CYP3RNA plants were sequenced and matching reads were aligned towards the Arabidopsis genome. (C) Sequenced siRNAs from 18-25 nt length from differential treated EVs were mapped towards the CYP3RNA precursor. Three siRNAs were found after PK and RA treatment, non was found after TX, PK and RA treatment. (D-F) RNA-seq data were grouped using featurecounts into different classes concerning their annotations as lncRNA, miRNA, mRNA, ncRNA, rRNA, snoRNA, snRNA and tRNA to analyse RNA compositions of different RNA origin. Reads which could not be annotated are declared as ‘not assigned’. (D) Comparison of RNA composition isolated from whole leaf cells and EVs purified by apoplastic washes of Arabidopsis wt plants. (E) Amount of total alignments of RNA isolated from *Δvps20*-CYP3RNA, *Δvps2*-CYP3RNA, *Δsnf7a*-CYP3RNA, *Δsnf7b*-CYP3RNA, and *Δsnf7c*-CYP3RNA EVs comparison to wt EV RNA. (F) Comparison of RNA species isolated from differential treated wt and CYP3RNA EVs.

Mapping of siRNA reads to the CYP3RNA precursor identified only three siRNAs matching to the CYP3RNA in protease and RNase treated EV samples while in the Triton X, protease and RNase treated EV samples no matching siRNAs were found (Figure 3C). Two of them mapped to the CYP51A fragment (5’-AGGTTCTTGACTACGTCGAAA-3’, 5’-TCCTTTTCTGGCAGAACTAGC-3’), one to CYP51B (5’-CCCCTGGAACCGTAAGCGC-3’) and none to CYP51C (Figure 3C).

Next, we analysed how digestive treatment influence RNA composition in general. For this, we mapped the RNA-seq data to the Arabidopsis genome and results were classified as long non-coding (lnc)RNA, micro (mi)RNA, messenger (m)RNA, non-coding (nc)RNA, ribosomal (r)RNA, small nucleolar (sno)RNA, small nuclear (sn)RNA and transfer (t)RNA based on most abundant annotations from TAIR10 and counted with featureCounts (Liao et al. 2019) to get an overview of RNA species composition. We compared the RNA species compositions from whole plants with the RNA composition of EVs isolated from apoplastic fluids after degradation of extravesicular RNAs by protease and RNase treatment and found a strong reduction of miRNA, ncRNA, snoRNA, snRNA and tRNA, while the mRNA amount was increased (Figure 3D). We then compared EV RNA composition after treatment with the EV-destructive detergent TX (prior to RNase and protease treatment) and found that ncRNAs showed no reduction after protease and RNase treatment but were reduced after additional TX treatment, while all other RNA species were constantly reduced by protease/RNase and triton/protease/RNase treatment (Figure 3F).

### ESCRT-III is required for HIGS-mediated *Fg* resistance

To further assess whether EVs are required for HIGS in Arabidopsis, we tested if impairment of the endosomal sorting complex, which is required for the transport III (ESCRT-III) pathway, affects HIGS-mediated *Fg* inhibition. We predicted that the translocation of transgene-derived siRNA to the fungus involves intact plant intraluminal vesicles (ILVs) and MVB-mediated release of EV at the plant-fungal interface (Koch and Kogel, 2014). To test this prediction, we first selected seven Arabidopsis ESCRT-III T-DNA KO mutants to assess whether a transfer of CYP3RNA-derived siRNAs is compromised in plants with a defective exosomal vesicle pathway. The mutants were co-transformed with the CYP3RNA construct and subsequently genotyped and propagated (Supplementary Table 4). The T2 generation of transgenic plants *Δvps2-*CYP3RNA, *Δvps20*-CYP3RNA, *Δrabf1*-CYP3RNA, *Δrabf2a*-CYP3RNA, *Δrabf2b*-CYP3RNA, *Δsnf7a*-CYP3RNA, *Δsnf7b*-CYP3RNA, and *Δsnf7c*-CYP3RNA were further investigated. Using a detached leaf assay, we found that virtually all tested CYP3RNA-generating ESCRT-III mutants were more susceptible to *Fg* as compared with CYP3RNA-expressing non-mutated plants (Figure 4A), hinting to the possibility that a functional ESCRT-III is required for HIGS.

**Figure 4.**
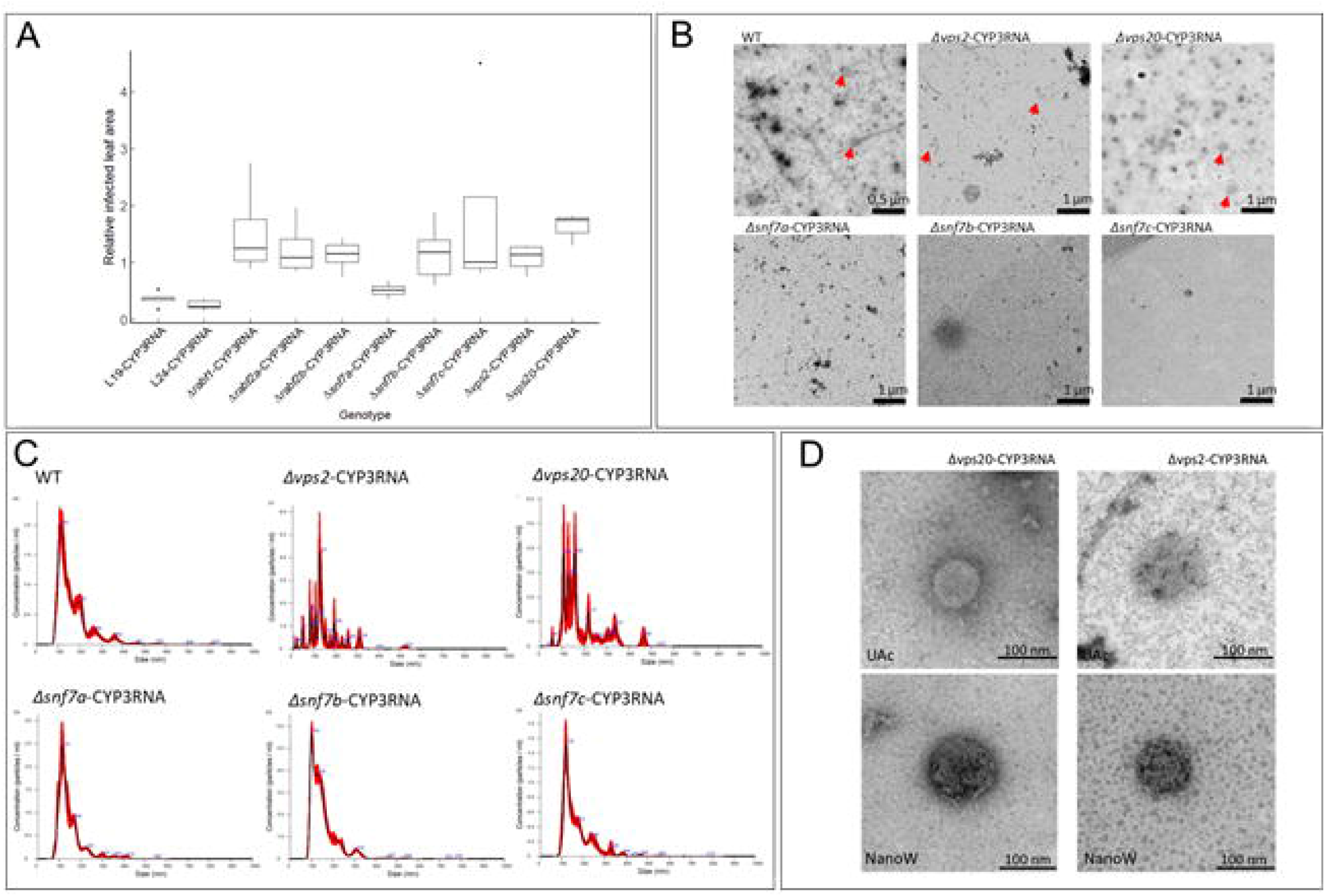
Analysis of ESCRT-III KO mutants in HIGS-mediated resistance to *Fg*. (a) HIGS in *Fg* upon infection of transgenic Arabidopsis expressing CYP3RNA in ESCRT-III knock-out mutant background. Fifteen detached rosette leaves of CYP51-dsRNA-expressing Arabidopsis plants (T2 generation) were drop-inoculated with 5 x 10^4^ conidia ml^-1^. Infection symptoms were quantified at 5 dpi and displayed as relative infected leaf area. The average relative infected leaf area was normalized against Col-0 control plants and a total of 3 to 6 repetitions were conducted, each using 15 leaves of 10 different plants for each transgenic line. Each dot represents one repetition and the underlying box and whisker plots represent the median and quartiles of said repetitions. Asterisks indicate statistical significance (**p<0,01; ***p<0,001; students t-test; two-sided). (b) EVs of wt, *Δvps20*-CYP3RNA, *Δvps2*-CYP3RNA, *Δsnf7a*-CYP3RNA, *Δsnf7b*-CYP3RNA, and *Δsnf7c*-CYP3RNA were isolated by apoplastic washes and prepared for TEM analysis by negative staining with UAc and NanoW. EVs or the single EV like structure of *Δsnf7a*-CYP3RNA were marked by red arrows. (c) Particle size measurements by NTA analysis of particles isolated by apoplastic washes from Arabidopsis wt, *Δvps20*-CYP3RNA, *Δvps2*-CYP3RNA, *Δsnf7a*-CYP3RNA, *Δsnf7b*-CYP3RNA, and *Δsnf7c*-CYP3RNA. (d) Mean and mode values of the particle diameter from Arabidopsis wt, *Δvps20*-CYP3RNA, *Δvps2*-CYP3RNA, *Δsnf7a*-CYP3RNA, *Δsnf7b*-CYP3RNA, and *Δsnf7co*-CYP3RNA isolated apoplastic particles. Diameters were calculated by the NTA 3.2 Dev Build 3.2.16 software for the NTA measurements seen in (c). (e) EVs were purified by apoplastic washes from Arabidopsis wt plants and Δ*vps2*-CYP3RNA and Δ*vps20*-CYP3RNA mutants. EVs were resuspended in PBS and stained with glutaraldehyde, formaldehyde and osmium tetraoxide before negative staining was performed by using 2 % uranyl acetate (UAc) and NanoW.

To substantiate this finding, we isolated EVs from the apoplastic washing fluid of the five mutant lines *Δvps20*-CYP3RNA, *Δvps2*-CYP3RNA, *Δsnf7a*-CYP3RNA, *Δsnf7b*-CYP3RNA, and *Δsn/7c*-CYP3RNA to analyse vesicle size distribution by TEM and NTA (Figure 4B, 4C). NTA revealed particle sizes between 100 nm and 150 nm in diameter (Figure 4C; Table 2). To further investigate the morphology of EVs by TEM, we stained lipids and proteins as well as fixed samples with uranyl acetate (UAc) and NanoW (Figure 4D). When EVs were stained with UAc, PBS buffer in the EV suspension leads to poor background signals. To obtain a better contrast EVs were negatively stained with NanoW. As expected, NanoW led to less background signals and higher contrast (Figure 4D). EVs from *Δvps20*-CYP3RNA were round shaped and had a smooth and clear boundary to the environment. However, UAc staining reveals a homogenous inner structure (Figure 4D), while NanoW staining showed heterogenous intravesicular structures like in wt EVs (Figure 4D). We have no explanation why both staining methods lead to such a different appearance. EVs of the *Δvps2*-CYP3RNA mutant instead have no clear shaped borders compared to wt and *Δvps20*-CYP3RNA plants independently of the staining agent (Figure 4D). The edges look very diffuse and rough while the inside has different and complex structures like EVs from the wt. For *Δsnf7a*-CYP3RNA, *Δsnf7b*-CYP3RNA, and *Δsnf7c*-CYP3RNA we observed no differences among EVs isolated from wt plants using NTA (Figure 4C). Important to note, TEM analysis revealed no EVs from *Δsnf7b*-CYP3RNA and *Δsnf7c*-CYP3RNA plants and only one EV-like structure in isolated EV samples of *Δsnf7a*-CYP3RNA (Figure 4B), indicating that co-isolated extracellular particles or aggregates led to similar/indistinguishable NTA results, which are only stable in liquid environments and therefore not visible during TEM, where samples are dehydrated during staining procedure. To substantiate our assumption that ESCRT-III-dependent EV transport is required for HIGS we performed sRNA-seq of EVs/apoplastic fluid isolated from *Δvps2-*CYP3RNA, *Δvps20*-CYP3RNA, *Δsnf7a*-CYP3RNA, *Δsnf7b*-CYP3RNA, and *Δsnf7co*-CYP3RNA transgenic plants. Of note, none of the mutant-EVs or mutant-derived apoplastic fluid contained CYP3RNA-derived siRNAs (Supplementary Figure 3). Moreover, analysing EVs’ RNA content revealed a decrease of all RNA species (Supplementary Figure 4).

**Table 2.** Average diameter of vesicles from Arabidopsis. Average diameter was analysed by NTA. Standard errors (SE) are given in parentheses. Arabidopsis CYP3, CYP3RNA-expressing plants.

| Plant | Vesicle diameter [nm] |
| --- | --- |
| Arabidopsis wt | Mean 167 (± 4) |
| | Mode 116 (± 4) |
| *At-Δvps2*-CYP3RNA | Mean 147 (± 14) |
| | Mode 114 (± 9) |
| *At-Δvps20*-CYP3RNA | Mean 182 (± 7) |
| | Mode 120 (± 7) |
| *At-Δsnf7a*-CYP3RNA | Mean 147 (± 2) |
| | Mode 116 (± 7) |
| *At-Δsnf7b*-CYP3RNA | Mean 155 (± 1) |
| | Mode 106 (± 5) |
| *At-Δsnf7c*-CYP3RNA | Mean 169 (± 2) |
| | Mode 119 (± 4) |

### Comparative proteomic analysis of Arabidopsis EVs

To gain further insight into the protein content of EVs we conducted a proteomic comparative analysis of Arabidopsis EVs using our data and compared them to published data (Rutter and Innes, 2017). To this end, EVs were isolated from apoplastic washes of 5-wk-old Arabidopsis wt plants as described (Rutter and Innes, 2017). By categorizing the proteins to GO-terms we observed a 2-fold enrichment of proteins associated with stress responses and abiotic stimuli compared to the whole proteome as previously described by Rutter and Innes (Rutter and Innes, 2017) (Figure 5A, 5B, 5C). Moreover, we observed a 6-fold and 2-fold enrichment of proteins associated with translation and proteins of the categories transport and cellular component organization, respectively. Comparing the identified proteins by Rutter and Innes (Rutter and Innes, 2017) with MS data of this study we found an overlap of 81 proteins identified in both studies (Figure 5D). Remarkably, these overlapping proteins are specifically proteins with high abundance in both studies and the 15 proteins with the highest abundance are identical. Following the procedure of Rutter and Innes (Rutter and Innes, 2017) we further classified the identified proteins to GO-terms via the PANTHER Classification System (Supplementary Figure 5A, 5B, 5C). However, we assume that differences in the protein profiles resulted from using plants of different age and divergent genotypes. For example, Rutter and Innes (Rutter and Innes, 2017) isolated EVs from 5-7-wk-old, genetically modified (GFP-PEN1) Arabidopsis plants, while we used 4-5-wk-old wt Col-0 Arabidopsis plants.

**Figure 5.**
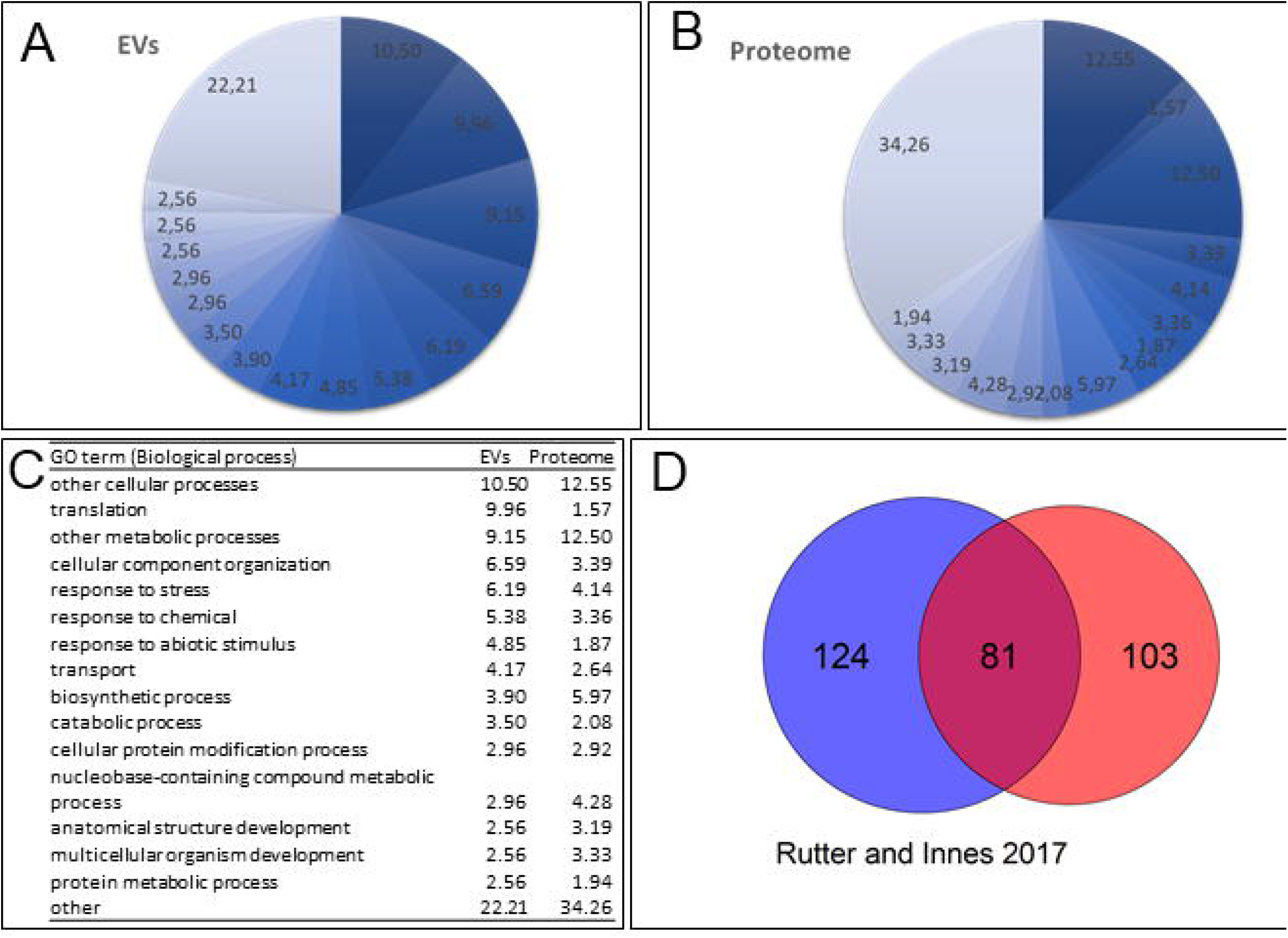
Functional categorization of EV-contained proteins. PSMs for UniprotKB accessions were linked to Arabidopsis genes with the R package biomaRt. UniprotKB accessions were searched in the athaliana_eg_gene dataset on plants.ensembl.org and linked to the corresponding ensembl_gene_id. These gene_ids were linked to Gene Ontology terms of the category biological processes. (a)(b)(c) The relative amount [%] of the 15 most abundant terms is shown for EV-contained proteins (a) compared to the whole proteome (b) with specific numbers and categories (c). (d) Venn-diagram comparing proteins found in EVs by Rutter and Innes [27] compared to the data obtained within this study. To accommodate for the higher number of PSMs in the later datasets only genes with at least 1‰ of total PSMs in the respective study were considered.

## Discussion

The discovery that plants have evolved the capability of silencing foreign transcripts trough RNAi by their own sRNAs (Zhang et al. 2016) or after biotechnological manipulation by sRNA derived from a dsRNA-generating transgene is especially relevant for future agricultural applications. The latter possibility, called HIGS (Nowara et al. 2010) has been used to successfully engineer resistance to a broad range of pests and microbial pathogens (Koch and Wassenegger 2021). However, one of the major gaps in knowledge is the transfer of transgene-expressed dsRNA, specifically dsRNA-derived siRNAs from the host plant cell to the recipient fungal target cell (Figure 6). Because fungi probably lack RNA transporters, such as SID1 and SID2 found in nematodes and insects, RNA uptake during plant-fungus communication via membrane-localized RNA transporters is an unlikely option. Thus, several recent reports suggest a bidirectional sRNA transport via EVs (Cai et al. 2018b, Rutter and Innes, 2018) in analogy to the role mammalian exosomes have in cell-to-cell communication (Maia et al. 2018, Mittelbrunn et al. 2011, Ratajczak et al. 2006, Takahashi et al. 2017, Valadi et al. 2007). In addition, sRNA cargo of EVs has been shown for fungi (Peres da Silva et al. 2015), nematodes (Quintana et al. 2017) and plant pathogenic bacteria (Katsir and Bahar, 2017). In mammalian cells, sRNAs of parasitic origin are transported to host cells via vesicles (Buck et al. 2014, Zhu et al. 2016). In line with this knowledge, we followed the hypothesis that sRNA transfer in HIGS also is vesicle-based. As an essential basis, we used developed protocols (Mu et al. 2014, Rutter and Innes, 2017) for the isolation of EVs and their cargo from Arabidopsis leaves. Vesicles isolated by these methods were around 140 nm (139 +/− 7.7 nm) in diameter (Figure 1; Figure 2), which is in good agreement with the size range reported for small EVs from mammalian cells (30-150 nm (Raposo and Stoorvogel, 2013)) and plants (50-300 nm (Rutter and Innes, 2017)). The physical appearance of vesicles in the TEM analysis was comparable to typical exosome preparations from cell culture supernatants (Li et al. 2017), especially as they were surrounded by a characteristic lipid bilayer which has an average thickness of around 5 nm in diameter (Figure 1; Figure 2). Moreover, mammalian EVs have a density of 1,13-1,19 g/ml (Szatanek et al. 2015), which fits well with the detection of the Arabidopsis vesicles in the sucrose gradient between the 30% and 45% sucrose corresponding to a density of 1.1270 g/ml and 1.2025 g/ml, respectively. Vesicles isolated from Arabidopsis cell extracts showed a slightly more cup-shaped morphology than those from the apoplastic fluid. Cup-shaping of exosomes is an effect of extreme dehydration during conventional TEM procedures (Li et al. 2017, Rutter and Innes, 2017). This effect can be more or less severe during different sample preparations, thus explaining the different appearance of Arabidopsis vesicles. Sucrose gradient centrifugation has been repeatedly used for mammalian EV purification and subsequent RNA sequencing. From our point of view, gradient centrifugation currently represents one of the most stringent method to obtain different EV subpopulations, according to their buoyant density (Alexander et al. 2016, van Balkom et al. 2015).

**Figure 6.**
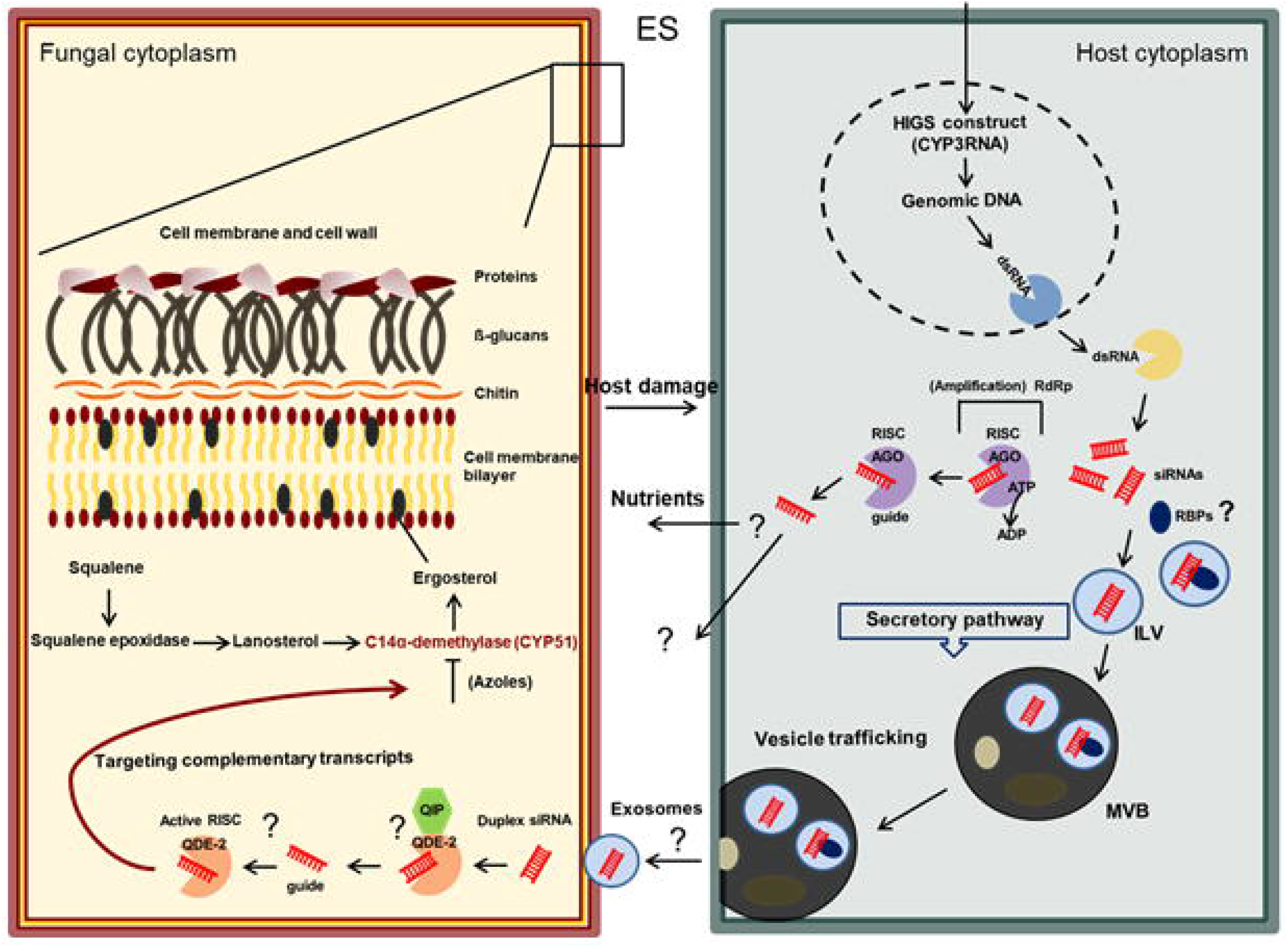
Vesicle-mediated transfer of HIGS-derived RNAs. A hypothetical siRNA translocation pathway involves the integration of a HIGS construct, its transcription into dsRNA and the translocation into the cytoplasm, where it is loaded and processed by DCL enzymes (yellow). The resulting siRNA duplexes are either delivered to the plant’s RNA silencing machinery or are incorporated as duplexes into intraluminal vesicles (ILVs) that originate either from the Golgi body via the trans-Golgi network or from endocytosis at the cell membrane, respectively (circles light blue). The ILVs, consisting of several cargos, are internalized by MVBs that enter the secretory pathway (circles dark grey). MVBs fuse to the PPM followed by subsequent release of ILVs (now called exosomes) (circles light blue). Exosomes cross the cellular interface, entering the fungal cell and release their cargo, possibly including plants siRNAs (process unknown). The siRNAs may subsequently enter the fungal RNAi machinery resulting in target gene silencing where they are wrenched by the AGO protein QDE-2 (quelling deficient-2), while the passenger strand is removed by the exonuclease QIP (hexagon green). The guide strand remains in the RNA-induced silencing complex (RISC) which is activated and targets complementary mRNAs, resulting in degradation and *CYP51* gene silencing, respectively.

Plant EV research is an emerging field, thus the methodology of EV isolation and preparation continuously undergoes further development and optimization. For example, leaf infiltration and harvest of the apoplastic fluid by depression seems to be a very harsh method that may cause symplastic contaminations through cell disruption. These artefacts may lead to mis- or overinterpretation of results. Given this assumption, we performed different treatments of EV isolates, i.e., protease and RNase digest to eliminate extravesicular proteins and RNAs as well as ribonucleoprotein complexes prior to RNA extraction and sequencing. Of note, stringent treatment of EV isolates revealed that most of the RNA was extravesicular (Figure 3). We found that the total amount of RNA and the total read number decreased drastically after protease and RNase treatments demonstrating only three CYP3RNA-derived siRNAs as intravesicular (Figure 3/Supplementary Table 1). Therefore, the central question is: How many or less siRNAs are required to induce HIGS? Moreover, are EVs required for HIGS or does the fungus takes up unprocessed/apoplastic/extracellular RNAs as shown for the SIGS-barley-*Fg* system (Koch et al. 2016).

Our results indicate that most CYP3RNA-derived siRNAs were present in the apoplast as well as their stabilization by RNA-binding proteins (decrease of RNA after protease/RNase digest compared to RNase digest alone). Formation of ribonucleoprotein complexes that resist RNase digestion is consistent with mammalian EV reports demonstrating protection of circulating miRNAs from plasma RNases (Arroyo et al. 2011). The question is whether these apoplastic non-EV-encapsulated/naked RNAs/proteins are procedural artefacts or whether this indicates a directed (active or passive) ESCRT-III-mediated secretion of HIGS-associated RNAs into the apoplast. The latter assumption raises the possibility for fungal uptake of apoplastic HIGS inducers instead of the requirement for an EV-mediated siRNA transfer. If they were secreted/released in the apoplastic space we need to reconsider working models and hypothesis that previously focussed on the uptake of intracellular siRNA mediated by EVs (Koch and Kogel 2014) (Figure 6).

Supporting the notion that MVBs are involved in host-pathogen communication via their sRNA cargos, our results indicate that the ESCRT-III is required for HIGS-mediated *Fg* disease resistance as we demonstrated that KO mutants of ESCRT-III lost their resistance towards *Fg* infection. To further verify our results, we harvested apoplastic fluids from ESCRT-III mutants and found EVs isolated from Δ*vps2*-CYP3RNA and Δ*vps20*-CYP3RNA plants by TEM and NTA analysis. Notably, both mutants produced and released EVs, which can be purified from apoplastic washes, what we did not anticipate at the beginning because of the findings from Cai et al (2014) where they showed that intraluminal vesicles are strongly reduced in the *vps2, vps20* and *snf7* mutants (Figure 4). We systematically measured diameters of wt Arabidopsis plants in comparison to both mutant lines, using NTA- and TEM-based analysis (Figures 4). Despite a slight size difference of Δ*vps2*-CYP3RNA, surface appearance and composition of both tested mutant lines were impaired supporting the importance of ESCRT-III components in CYP3RNA-mediated *Fg* resistance. *Δvps20* and *Δvps2* mutants are compromised in EV functionality as VPS20 (VACUOLAR PROTEIN SORTING-ASSOCIATED PROTEIN 20) is recruited by ESCRT-II to start the assembling of ESCRT-III, while VPS2 is needed for EV membrane invagination and the formation of EVs. Thus, VPS20 functions as linker between ESCRT-I or ESCRT-II to ESCRT-III (Henne et al. 2013). Therefore, we anticipate that *vps20* mutants were impaired in correct protein/RNA packaging into EVs leading to the release of CYP3RNA-free EVs. VPS2 is recruited by SNF7 and forms oligomers which leads to sculpting the MVB membrane for forming a dome which then forms an EV inside MVB (Henne et al. 2013). The RAB (RAS-REALATED-PROTEIN) GTPases are key players in the regulatory process of endosomal vesicle transport. Apoplastic fluids of *Δsnf7a*-CYP3RNA, *Δsnf7b*-CYP3RNA, and *Δsnf7c*-CYP3RNA plants contain particles of same size and concentration during NTA analysis, but we found no EVs during TEM. These diverging results might be due to hydro stable complexes, which were lost after dehydration during TEM preparation. The same effect could cause the size differences measured between NTA and TEM, which displays slightly bigger particle sizes during NTA than EV size measurement by TEM for wt, Δ*vps2*-CYP3RNA and Δ*vps20*-CYP3RNA plants. Moreover, it shows the contamination with co-purified particles and complexes underlining the requirement of a more sensitive/stringent EV isolation method.

RNA-seq analysis of apoplastic fluids from all tested ESCRT-III mutants revealed no CYP3RNA-derived siRNAs, which may explain higher sensitivity towards *Fg* infection and the requirement of a functional EV biogenesis in HIGS-mediated *Fg* resistance. Of note, consistent with our data, Cai et al. (2018b) demonstrated that MVBs fused to the plasma membrane releasing EV-packaged plant sRNAs at fungal infection sites to silence virulence genes in *Botrytis cinerea* thus suppressing fungal pathogenicity (Cai et al. 2018b). This finding is supported by several lines of evidence from earlier reports: Fungal and bacterial infection activated the biogenesis and enhanced the formation of vesicles between cell wall and cell membrane, and these vesicles contain defence related compounds (An et al. 2006b, Rutter and Innes, 2017, Wang et al. 2014). Arabidopsis LIP5 (HOMOLOG OF MAMMALIAN LYST-INTERACTING PROTEIN5), a positive regulator of MVB biogenesis, is a critical target of pathogen-responsive MAPK cascade in plant basal defence (Wang et al. 2014). MVBs and paramural bodies proliferate underneath papillae in barley cells during powdery mildew infection. MVBs were also observed in fungal haustoria and in association with plasmodesmata, suggesting that the secretion of EVs contributes to the formation of defensive appositions, but may also help to block plasmodesmata in cells undergoing HR and contribute to defense-related endocytosis (An et al. 2006a, Micali et al. 2011). The PEN1 protein is secreted outside of the cell during powdery mildew infection and is encased in the papillae along with membranous material. The same is true for PEN3, SNAP33 (SOLUBLE N-ETHYLMALEIMIDE-SENSITIVE FACTOR ADAPTOR PROTEIN 33), and VAMP722 (VESICLE-ASSOCIATED MEMBRANE PROTEIN 722), suggesting that PEN1 is secreted inside vesicles (Figure 2D), like exosome secretion in mammals (Meyer et al. 2009). Finally, EVs isolated from the apoplastic fluids of sunflower (*Helianthus annuus*) contained families of proteins commonly found in mammalian and Arabidopsis EVs. EVs labeled with FM 4-64 were taken up by *Sclerotinia sclerotiorum* spores, reduced hyphal growth and permeabilized fungal membranes, as evidenced by the uptake of otherwise non-permeable dyes (Regente et al. 2017). However, given the low amounts of siRNAs found in EVs, which raised the question on physiological activities of those siRNAs for HIGS, we hypothesise that the ESCRT-III function in secretion of dsRNA-derived siRNAs into the apoplastic space. Therefore, we suggest that ESCRT-III’s role in HIGS does not solely rely on the release of EVs or EV-contained HIGS inducers.

In addition to RNase and protease digest we treated EV isolates with an EV-disruptive detergent TritonX (TX) as a control. We hypothesized to find no CYP3RNA-derived siRNAs in EVs upon EV disruption and protease/RNase digest. Indeed, we found no matches to the CYP3RNA precursor as expected (Figure 3C). Even more important, analysis of RNA species composition revealed a mixture of all RNA types intra- and extravesicular revealing apoplastic contamination and misinterpretation of EV isolated components. Only lncRNAs and snoRNAs were completely degraded after TX treatment (Figure 3F). Based on these findings, we speculate that probably not all types of RNA respond to/be affected by RNase digest. We treated vesicles with RNase A, an endoribonuclease that specifically degrades RNA at C and U residues, to further remove potential contamination from outside adhering RNA. Based on our results we suggest that a cocktail/mixture of different RNase is needed to ensure complete degradation of copurified RNA contaminants. However, whether all found RNA species are functional or degradation products on their way to disposal remains unclear. The many pitfalls in plant EV research were recently reviewed by Rutter and Innes (2020) exemplifying the requirement of EV markers, EV reference proteins/RNAs to ensure comparability, reliability, and reproducibility among different EV preparations. Thus, the central question is how robust are results from one EV preparation and sequencing event to another?

Moreover, we need to clarify the contribution of host and target RNAi in a pathosystem-specific manner. Based on sRNA-seq data that reveal 5’-identities and lengths of HIGS-derived siRNAs, we can speculate about DCLs and AGOs involved in processing of transgene-derived dsRNA and binding of siRNAs/loading onto RISC. Currently, we assume that HIGS relies on the plant’s silencing machinery (DCL processing), but we lack experimental evidence for this assumption. For SIGS we found clear indications that processing of long sprayed dsRNA by fungal DCLs is required for SIGS-mediated disease resistance (Koch et al. 2016; Gaffar et al. 2019). Given the uptake of unprocessed dsRNA from the apoplast we assume that this may explain higher SIGS efficiencies compared to HIGS as observed (Koch et al. 2019; Höfle et al. 2019). In line with this and together with the results we obtained within this study, we predict that RNAi-based plant protection strongly depends on the lifestyle of the target pathogen and pest. The question is whether biotrophs and necrotrophs acquire the same set of HIGS inducers depending on their divergent lifestyles. In addition, we lack information on the composition of HIGS inducers in the intra-, inter-, and extracellular space as well as the threshold of siRNAs required for induction.

Here we provided significant insights into RNA uptake, processing and transport that is a step forward in unrevealing the mechanistic basis of HIGS. Together our findings support the view that HIGS includes processing of dsRNA in the plant, incorporation of the produced siRNA into intracellular vesicles, secretion of the vesicles into the apoplastic space, and a potential uptake of the vesicles or their cargo into the interacting fungus (Figure 6). Although, it is an exciting possibility that plant EVs mediate the interspecies transfer of RNAs in natural crosskingdom communications (Cai et al. 2018a, Rutter and Innes, 2018), whether this holds true for HIGS requires further research. Our results clearly show the requirement for fundamental knowledge on the molecular mechanisms (i.e., uptake, processing, and translocation of transgene-expressed double-stranded RNAs) that determine the efficacy and specificity of HIGS necessary to further develop RNAi-based plant protection technologies.

## Material and Methods

### Isolation of vesicles from Arabidopsis leaves

For the isolation of vesicles, a published protocol was adjusted (Mu et al. 2014). Ten 4-5-wk-old Arabidopsis rosette leaves were harvested and ground with pestle and mortar in a small amount of 1x PBS buffer (140 mM NaCl, 2.5 mM KCl, 10 mM Na2HPO4, 2 mM KH2PO4, pH 7.4). The cell extract was collected on ice and larger particles were separated by sequentially centrifuged at 1,000 x*g* for 10 min (Eppendorf 5810R), 3,000 x*g* for 20 min, and 10,000 x*g* for 40 min at 4°C (Beckmann J2-21M/E). After the last centrifugation step, the supernatant was centrifuged at 160,000 x*g* for 90 min in an ultracentrifuge (Beckmann SW 41 Ti). The pellet was resuspended in 1 ml 1x PBS, loaded on top of a sucrose discontinuous gradient (8% / 15% / 30% / 45% / 60% w/v in 20 mM HEPES, pH 7.3), and centrifuged at 160,000 x*g* for 2 h. The bands between the 30% / 45% and the 45% / 60% sucrose layer were harvested separately. The respective fraction was filled up to 10 ml with 1x PBS and centrifuged again for 2 h at 160,000 x*g*. The pelleted vesicles were resuspended in a small amount of 1x PBS and analysed by TEM or subjected to RNA extraction and RNA sequencing.

### Isolation of EVs from apoplastic washes of Arabidopsis leaves

A published protocol was adjusted to isolate EVs from Arabidopsis leaves by apoplastic washes (Rutter and Innes, 2017). Briefly, 5-wk-old rosette leaves were harvested, and vacuum infiltrated with vesicle isolation buffer (VIB: 20 mM MES, 2 mM CaCl2 and 0.1 M NaCl, pH 6) until infiltration sites were visible. The infiltrated leaves were drained carefully on filter paper and centrifuged (Eppendorf 5810R) in 30 ml syringes placed in 50 ml falcon tubes for 20 min at 700 *g* and 4°C. The resulting apoplastic fluid was filtered through 0.45 μm sterile filters, followed by centrifugation at 10,000 *g* for 30 min at 4°C (Eppendorf 5417R). The supernatant was then diluted to 10 ml with VIB and centrifuged for 1 h at 48,000 *g* at 4°C (Beckmann J2-21M/E). The resulting pellet was washed with 10 ml PBS and finally resuspended in a small amount of PBS. For further analysis samples were stored at 4°C for a maximum of two days.

### Negative staining and transmission electron microscopy (TEM)

For TEM, copper formvar-coated 300-mesh electron microscopy grids were glow discharged prior to sample application for 40 sec. Subsequently, 5 μl of the sample resuspended in PBS were applied to the grids. Samples were dabbed off using Whatman filter paper and grids were washed three times in 50 μl of 2% uranyl acetate. Additionally, protein and lipid staining were performed. 55 μl vesicles were incubated for 3 min with 0.1% solution of glutaraldehyde and 3% paraformaldehyde for protein staining. Afterwards, vesicles were washed with PBS before samples were incubated with 1% osmiumtetraoxide solution for 1 min for lipid staining, followed by a second washing step with PBS. All dyes were diluted in PBS before staining. Finally, samples were negative stained and fixed with 2% uranyl acetate or NanoW, an organotungsten based negative staining agent, as described before and washed with water. Needless staining or fixing solutions, buffers and water were removed by Whatman paper between each step. Finally, grids were air dried. Preparations were inspected at 120 kV under zero-loss conditions (ZEISS EM912a/b) and images were recorded at slight underfocus using a cooled 2k x 2k slow-scan ccd camera (SharpEye / TRS) and the iTEM software package (Olympus-SIS).

### Vesicle size and concentration measurements by nanoparticle trafficking analysis (NTA)

For size and concentration prediction, purified EVs of Arabidopsis (wt), CYP3RNA, Δ*vps2*-CYP3RNA, Δ*vps20*-CYP3RNA, *Δsnf7a*-CYP3RNA, *Δsnf7b*-CYP3RNA, and *Δsnf7c*-CYP3RNA were diluted (1:50) with PBS. Subsequently, 500 μL of vesicle suspension was loaded into Nanosight NS300 (Malvern Panalytical). 5 measurements were performed at 25°C and size, concentration prediction and statistical analysis were performed by the NTA 3.2 Dev Build 3.2.16 software.

### PEN1 detection by immunoblot analysis

EVs were treated by 1% Triton100 water solution (Sigma-Aldrich), 4 μl of 10 μg ml^-1^ trypsin in 10 mM HCl or both beside an untreated control (Rutter and Innes, 2017). Protein of whole cell extract and of EVs were separated by electrophoresis and blotted onto a nitrocellulose membrane. The membrane was blocked by 5% milk powder in Tris buffered saline (TBS; 100 mM Tris, 150 mM NaCl) containing 0.1 % Tween at 4°C over night. After blocking the membrane was incubated with an anti-PEN1 antibody (Rutter and Innes, 2017) 1:5000 for 4 h at RT. A secondary antibody (Thermo Fisher Scientific) linked to horseradish peroxidase was diluted 1:10000 in TBS and used for one hour at RT before detection by chemiluminescence using Amersham^TM^ ECL^TM^ Prime Western Blotting Detection Reagent (GE Healthcare) following manufactures instructions.

### EV treatments and RNA extraction for RNA sequencing

Typically, vesicles were treated with RNase (0.4 ng μl^-1^ RNase A; Thermo Fisher Scientific) to remove extravesicular RNA before RNA isolation. Samples were incubated for 10 min at 37°C and then kept on ice. To determine RNA localisation to intra- or extravesicular space EVs of wt and CYP3RNA plants from the same isolation were divided into three groups. The first untreated group contains all RNAs which were intra- or co-purified extravesicular. The second group was treated with proteinase (1 ng μL^-1^ Proteinase K Thermo Fisher Scientific) and RNase (0.4 ng μl^-1^ RNase A; Thermo Fisher Scientific) to degrade extravesicular proteins, RNAs and more stable ribonuclear complexes. In the third group, EVs were destroyed by Triton X and subsequently intra- and extravesicular content was degraded by proteinase and RNase treatment. During the degradation process, all three groups were kept at 37°C for 30 min. Vesicle RNA was isolated using the Single Cell RNA Purification Kit (Norgen Biotek) according to the manufacturer’s instructions described for cells growing in suspension. RNA concentrations were determined using the NanoDrop spectrophotometer (Thermo Fisher Scientific) and RNA was stored at −80°C.

### Determine siRNAs originating from CYP3RNA

Indexed sRNA libraries were constructed from RNA isolated from vesicles with the TruSeq® Small RNA Library Prep Kit (Illumina) according to the manufacturer’s instructions. Indexed sRNA libraries were pooled and sequenced on the Illumina MiSeq Platform (1 x 36 SE) and the sequences were sorted into individual datasets based on the unique indices of each sRNA library. The quality of the datasets was examined with FastQC before and after trimming (https://www.bioinformatics.babraham.ac.uk/projects/fastqc/). The adapters were trimmed using cutadapt (Martin, 2011) version 2.8. To filter out bacterial contaminations kraken2 Wood, 2019) version 2.1.1 was used with the database obtained from the MGX metagenomics application (Jaenicke, 2018). All reads marked as unclassified were considered to be of non-bacterial origin and used for the subsequent alignment. The trimmed and filtered reads were mapped to the CYP3RNA sequence using bowtie2 (Langmead and Salzberg, 2012) version 2.3.2. to identify siRNAs derived from the precursor dsRNA sequence. The mappings were first converted into bedgraph using bedtools (Quinlan and Hall, 2010) version 2.26.0 and then to bigwig using bedGraphToBigWig (Kent et al. 2010). These files were used for visualization with IGV (Thorvaldsdottir et al. 2013). Read coverage is defined as the number of reads that match at a certain position of the sequence.

### Determine frequency of different RNA species

To determine RNA species, reference genome and annotation of Arabidopsis (TAIR10) were downloaded from EnsemblPlants (Howe et al. 2020). Adapter trimming of raw reads was done with TrimGalore (https://www.bioinformatics.babraham.ac.uk/projects/trim_galore/) (version 0.6.4 which used cutadapt (Martin, 2011) version 2.8. In this process all reads which became shorter than 18 nt were filtered out. Afterwards, nucleotides with a phred score below 20 and reads retaining less than 90% of their nucleotides in this process were removed using FASTQ Quality Filter from the FASTX-toolkit (https://github.com/agordon/fastx_toolkit) version 0.0.14. The bacterial contaminations were filtered out as demonstrated in the previous section. The remaining reads were aligned to the reference genome using STAR (Dobin, 2013) version 2.7.3a. The number of different RNA species was examined in R (R Core Team, 2020) version 4.0.2 using featureCounts from the package Rsubread (Liao et al. 2019) version 2.2.5 .featureCounts was run for each sample using the previously downloaded annotations of Arabidopsis. Following RNA types were examined: “lncRNA”, “miRNA”, x“mRNA”, “ncRNA”, “rRNA”, “snoRNA”, “snRNA” and “tRNA”. All alignments that could not be assigned to a feature were considered “not assigned”.

### Generation of transgenic Arabidopsis plants

The ESCRT-III knock-out (KO) mutants NASC ID: *vps2*-N661671, *vps20*-N669857, *snf7a*-N547108, *snf7b*-N676240, *snf7c*-N674613, *rabf1*-N658064, *rabf2a*-N512028, *rabf2b*-N656748 were obtained from Nottingham Arabidopsis Stock Centre (NASC) (Supplementary Table 3) and transformed to express CYP3RNA (as described in Koch et al. 2013) in ESCRT-III KO background resulting in the following transgenic lines: Δ*vps2*-CYP3RNA, Δ*vps20*-CYP3RNA, Δ*snf7a*-CYP3RNA, Δ*snf7b*-CYP3RNA, Δ*snf7c*-CYP3RNA, Δ*rabf1*-CYP3RNA, Δ*rabf2a-CYP3RNA* and Δ*rabf2b*-CYP3RNA. The plasmid was introduced into *Agrobacterium tumefaciens* strain AGL1 (Lazo et al. 1991) by electroporation. Transformation of Arabidopsis was performed as described (Bechtold and Pelletier 1998) and transgenic plants were selected on 1/2 x MS (Murashige and Skoog) agar plates containing 7 μg ml^-1^ BASTA (Duchefa). The transgenic lines for pathogen assays were selected based on transgene expression levels measured by qRT-PCR.

### Plant infection assays

*Fusarium graminearum* (*Fg*) strain *Fg*-IFA65 (IFA, Department of Agrobiotechnology, Tulln, Austria was cultured on synthetic nutrient-poor agar (SNA)-medium (Gaffar and Koch, 2019). Plates were incubated at room temperature (RT) under constant illumination from one near-UV tube (Phillips TLD 36 W/08) and one white-light tube (Phillips TLD 36 W/830HF). For all leaf inoculation assays, *Fg* conidia suspensions were adjusted to 5 × 10^4^ conidia per ml^-1^ (Koch et al. 2013). *Arabidopsis thaliana* Col-0 wild type (wt) and transgenic plants were grown in a climate chamber under short day conditions with 8 h light at 22°C with 60% relative humidity. For infection, twenty rosette leaves of 15 5-wk-old Arabidopsis plants were detached and transferred in square Petri plates containing 1% water agar. Detached leaves were wound inoculated as described (Koch et al. 2013). Lesion size (in millimeters) was recorded 3 days post inoculation (dpi) from the digital images using the free software ImageJ program (http://rsb.info.nih.gov/ij/index.html). The percentage of leaf area showing water-soaked spots with chlorotic and/or necrotic lesions (symptoms of a successful *Fg* infection; (Koch et al. 2013)) relative to the non-inoculated leaf was calculated.

### In-solution tryptic digestion

Pellets of isolated EV fractions were washed twice with 50 mM ammonium bicarbonate buffer using Amicon Ultra 3 kDa cut-off centrifugal filters (Merck Millipore Ltd), and total protein concentration was estimated using Quick Start Bradford Protein Assay (Bio-Rad Laboratories). Protein samples were subjected to in-solution tryptic digestion followed by mass spectrometry (MS) analysis as described earlier with minor modifications (Vannuruswamy et al. 2017). In brief, proteins (20 μg) were diluted with 50 mM ammonium bicarbonate buffer containing 0.1% RapiGest (Waters Corporation), followed by incubation at 80°C for 15 min. The denatured proteins were then reduced with 100 mM dithiothreitol for 15 min at 65°C, and alkylated with 200 mM iodoacetamide at room temperature in the dark for 30 min. Sequencing grade modified trypsin (Promega Corporation) was added in 1:20 ratio (trypsin/protein, w/w) and the mixture was incubated overnight (16 h) at 37°C. Tryptic digestion reaction was stopped by adding 2 μl of concentrated formic acid and incubated for 10 min at 37°C before being centrifuged. Digested peptides were desalted using C18 ZipTips (Millipore) and concentrated using vacuum concentrator. Dried peptides were reconstituted in aqueous 3% acetonitrile with 0.1% formic acid.

### Proteomic analysis

Peptide digests (2 μg) were separated by using a Dionex Ultimate 3000 UHPLC system (Thermo Fisher Scientific) equipped with a reversed-phase Kinetex C18 2.6 μm column (100 X 2.1 mm, 100 A°). The sample was loaded onto the column with 97% of mobile phase A (100% water with 0.1% formic acid) and 3% mobile phase B (100% acetonitrile with 0.1% formic acid) at 250 μl/min flow rate. Peptides were eluted with a 120 min linear gradient of 3 to 50% of mobile phase B. The column and autosampler compartments were maintained at 40°C and 4°C, respectively. Proteomic datasets were acquired in a data-dependent (ddMS^2^-top 10) manner using a hybrid quadruple orbital trapping mass spectrometer (Q Exactive; Thermo Fisher Scientific) in positive-ion mode as described earlier (Korwar et al. 2015). MS data files (three biological replicates) were processed using Proteome Discoverer software (version 2.2.0.388, Thermo Fisher Scientific) with Sequest HT search algorithm. The raw data were searched against UniProt Arabidopsis protein datasets (reviewed, downloaded on 19^th^ June 2018). Search parameters included trypsin as proteolytic enzyme with two missed cleavages and 1% false discovery rate. Peptide precursor and fragment mass tolerance were set to 10 ppm and 0.6 Da, respectively. Search criteria also included carbamidomethylation (C) as fixed modification, protein N-terminal acetylation and oxidation (M) as variable modifications. As regulated, only proteins with a minimum of two peptides were considered for further analysis. Identified proteins were categorized according to molecular functions, biological processes, and cellular components based on gene ontology (GO) annotations using “The Arabidopsis Information Resource” (https://www.arabidopsis.org/tools/bulk/go/index.jsp).

## Acknowledgements

We thank C. Birkenstock, U. Schnepp and V. Weisel for excellent plant cultivation and M.Sc. C. Pfafenrot and M.Sc. M. Mosbach for helping with the NTA measurements. We thank Roger Innes for providing us the PEN1 antibody. We thank the Salk Institute Genomic Analysis Laboratory for providing the sequence-indexed Arabidopsis T-DNA insertion mutants. This work was supported by the Deutsche Forschungsgemeinschaft, Research Training Group (RTG) 2355 (project number 325443116) to AK and TS. VG thanks the German Academic Exchange Service (DAAD) for a doctoral fellowship. Financial support by the Deutsche Forschungsgemeinschaft (Sp 314/13-1 and INST 162/500-1 FUGG) is gratefully acknowledged. We acknowledge access to compute resources of the Bielefeld-Gießen Center for Microbial Bioinformatics (BiGi) financially supported by the BMBF grant FKZ 031A533 within the de.NBI network.

## Author Contributions

T.S., L.W. and A.K, wrote the manuscript; A.K. and T.S. designed the study; T.S., L.W., D.B., V.G., and M.C. conducted the experiments; A.K., T.S., L.W., V.G., B.S., P.B. and B.W. analyzed all data and drafted the figures. L.W. and T.B. conducted RNA-seq experiments and B.W., T.B., J.K., P.B. and L.J. performed bioinformatics analysis. All authors reviewed the final manuscript.

## Supplementary figures

**Supplementary Figure 1.** TEM analysis of fraction F160-s2 (A) and the interphase fraction (B) between the visible bands corresponding to F160-s1 and F160-s2 of the sucrose gradient shown in Figure 1A. Material was prepared from CYP3RNA-expressing Arabidopsis rosette leaves.

**Supplementary Figure 2**. sRNA-seq Col-0 (sucrose gradient). sRNA profiling of EVs isolated from leaf cell extracts from CYP3RNA-expressing (CYP3RNA-whole leaves) and wt (wt-whole leaves) Arabidopsis plants. Tracks show the read coverage of sRNAs on the CYP3RNA construct (logarithmic scale, 0-10 reads). There are no reads matching the CYP3RNA in wt plants (A). sRNA-seq Col-0 (EVs). sRNA profiling of apoplastic EVs isolated from CYP3RNA-expressing (CYP3RNA-apoplast) and wt (wt-apoplast) Arabidopsis plants. Tracks show the read coverage of sRNAs on the CYP3RNA construct (logarithmic scale, 0-10 reads). There are no reads matching the CYP3RNA in wt plants (A). NTA size distribution of particles isolated by apoplastic washes from Arabidopsis wt plants (B).

**Supplementary Figure 3.** siRNA profile of EVs purified by apoplastic washes and treated with RNase from Arabidopsis wt, *Δvps20*-CYP3RNA, *Δvps2*-CYP3RNA, *Δsnf7a-CYP3RNA, Δsnf7b*-CYP3RNA, and *Δsnf7c*-CYP3RNA and CYP3RNA plants. Only RNAsey of CYP3RNA EVs revealed matching siRNAs.

**Supplementary Figure 4.** RNA species compositions of of RNA isolated by apoplastic washes from wt, *Δvps20*-CYP3RNA, *Δvps2*-CYP3RNA, *Δsnf7a*-CYP3RNA, *Δsnf7b*-CYP3RNA, and *Δsnf7c*-CYP3RNA plants.

**Supplementary Figure 5.** (A) Bioinformatics analysis (Functional categorization based on GO annotation) of the Arabidopsis Col-0 EV proteome dataset depicting the biological processes involved by the proteins. (B) Bioinformatics analysis (Functional categorization based on GO annotation) of the Arabidopsis Col-0 EV proteome dataset depicting the cellular components of the proteins. (C) Bioinformatics analysis (Functional categorization based on GO annotation) of the Arabidopsis Col-0 EV proteome dataset depicting the molecular functions of the proteins.

**Supplementary Table 1.**
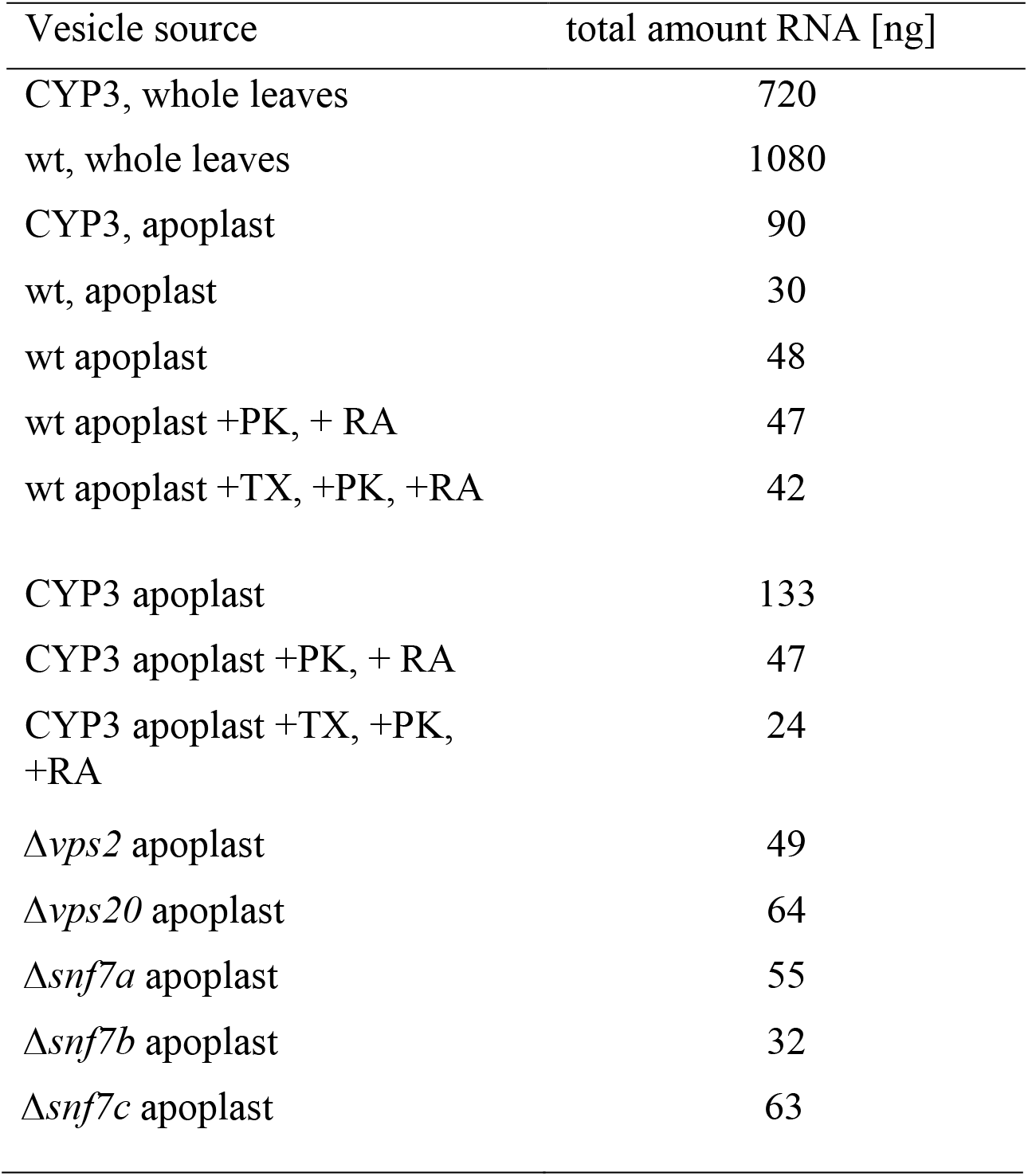
Amount of RNA isolated from different vesicle preparations used for cDNA library preparation and subsequent RNA sequencing. At-CYP3: CYP3RNA-expressing Arabidopsis, wt: wild type.

**Supplementary Table 2.** Number of reads (million) in datasets from sRNA sequencing of plant vesicle sRNAs. Read number in million is shown before and after adapter trimming. Vesicles from CYP3RNA-expressing Arabidopsis (*At*) leaves.

| Vesicle source | No. of reads | No. of reads after trimming |
| --- | --- | --- |
| *At*-CYP3, whole leaves | 5.7 | 5.3 |
| *At*-wt, whole leaves | 2.3 | 2.1 |
| *At*-CYP3, apoplast | 5.4 | 2.4 |
| *At*-wt apoplast | 2.9 | 0.7 |

**Supplementary Table 4.**
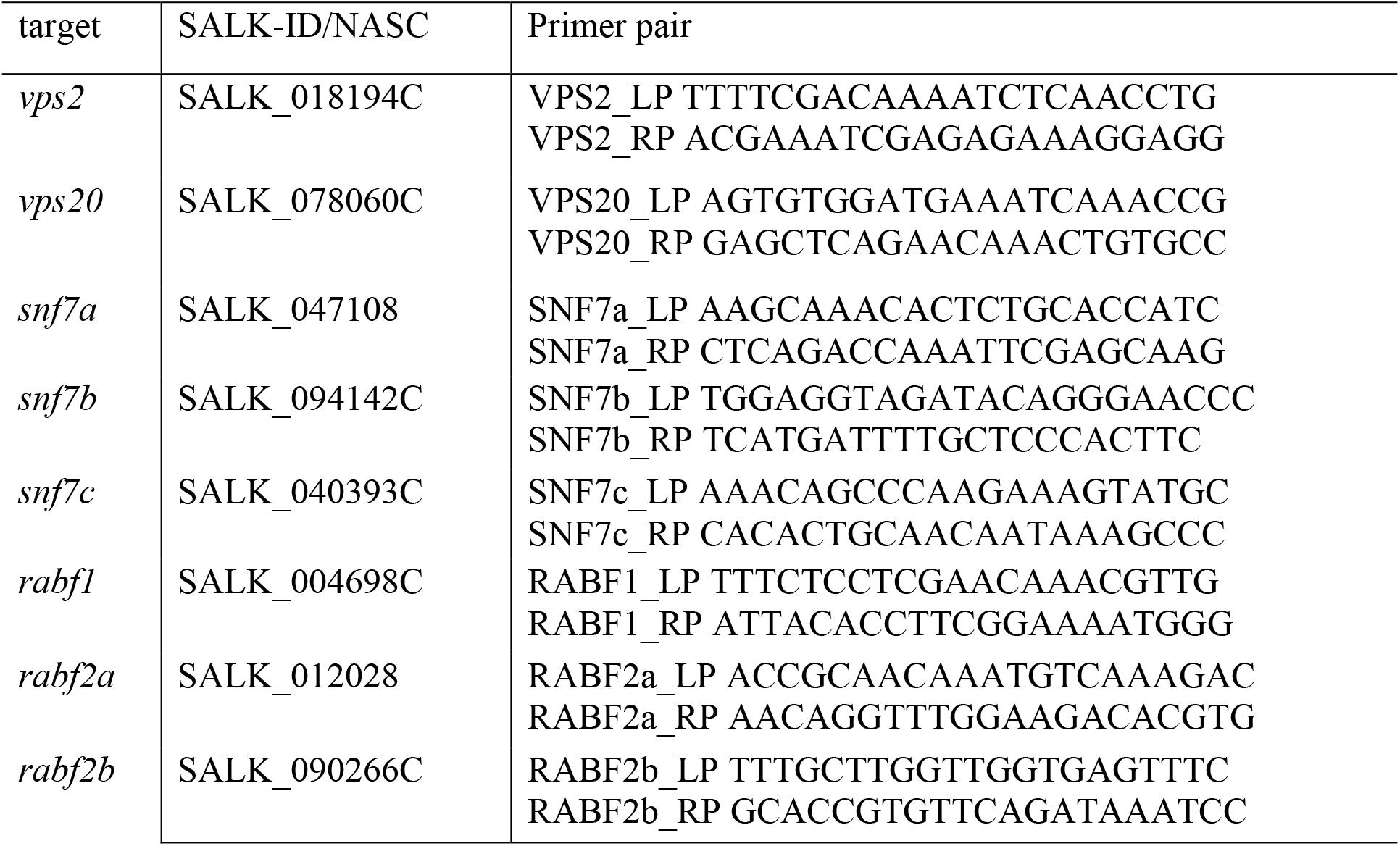
List of primers used for genotyping of ESCRT-III KO mutants

