## Supplementary figures and images for "Host-induced gene silencing involves Arabidopsis ESCRT-III pathway for the transfer of dsRNA-derived siRNA"

### Fig S1

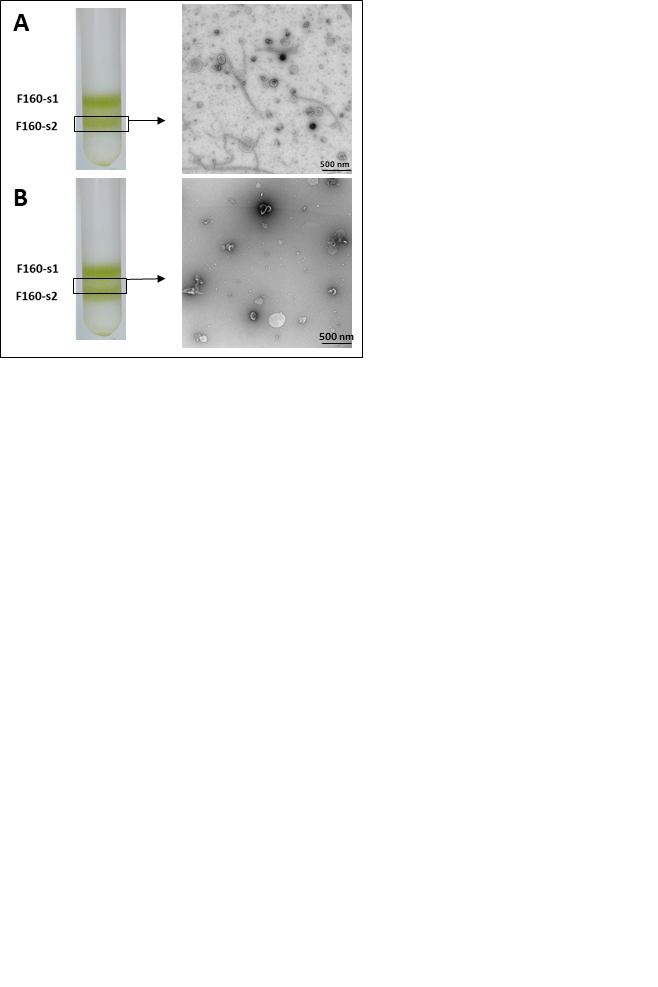

### Fig S2

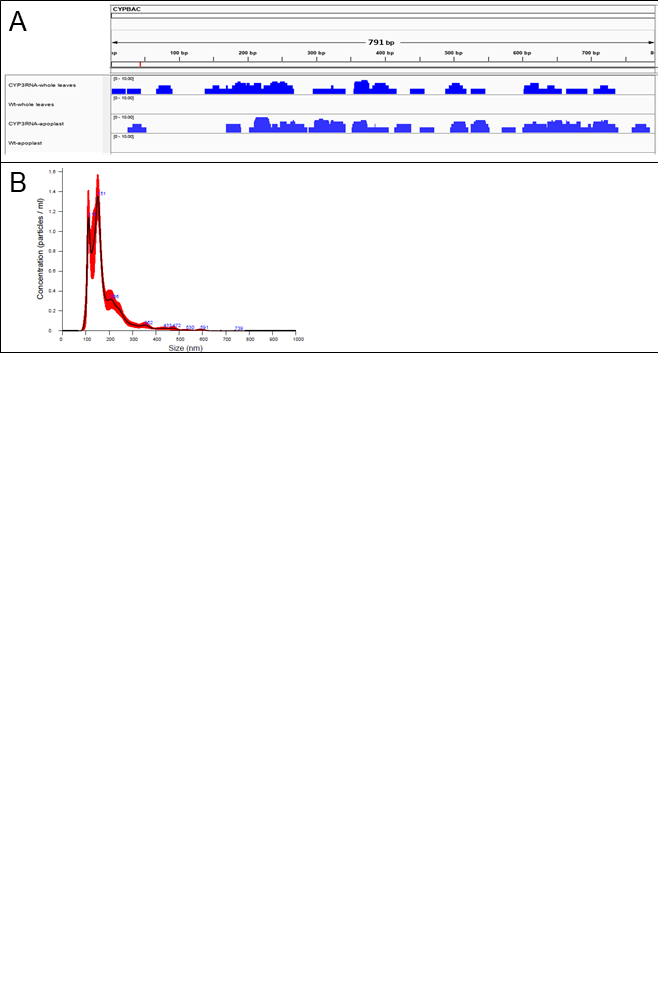

### Fig S3

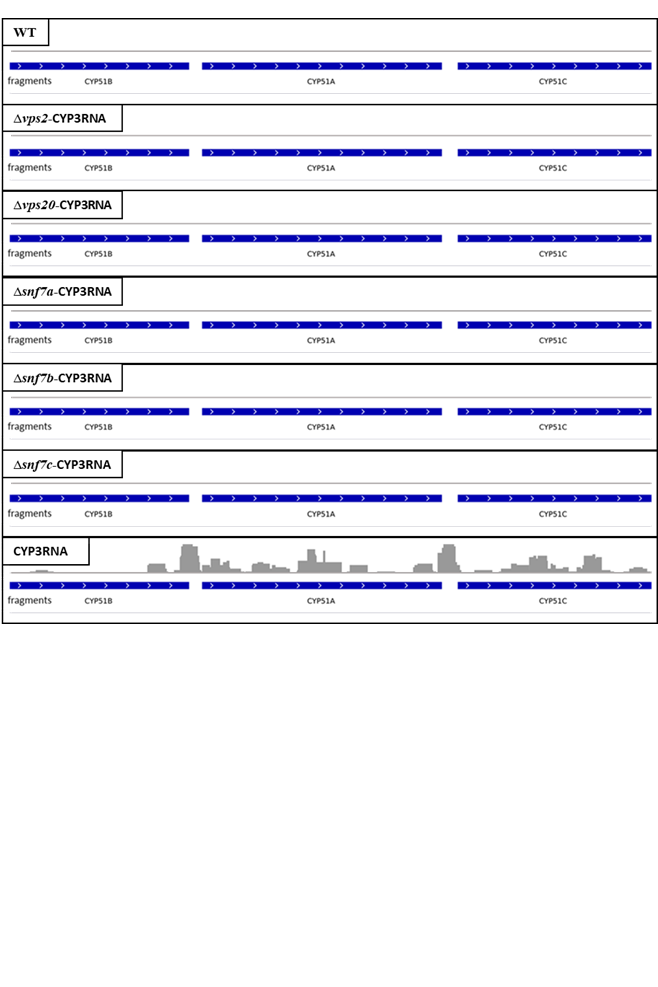

### Fig S4

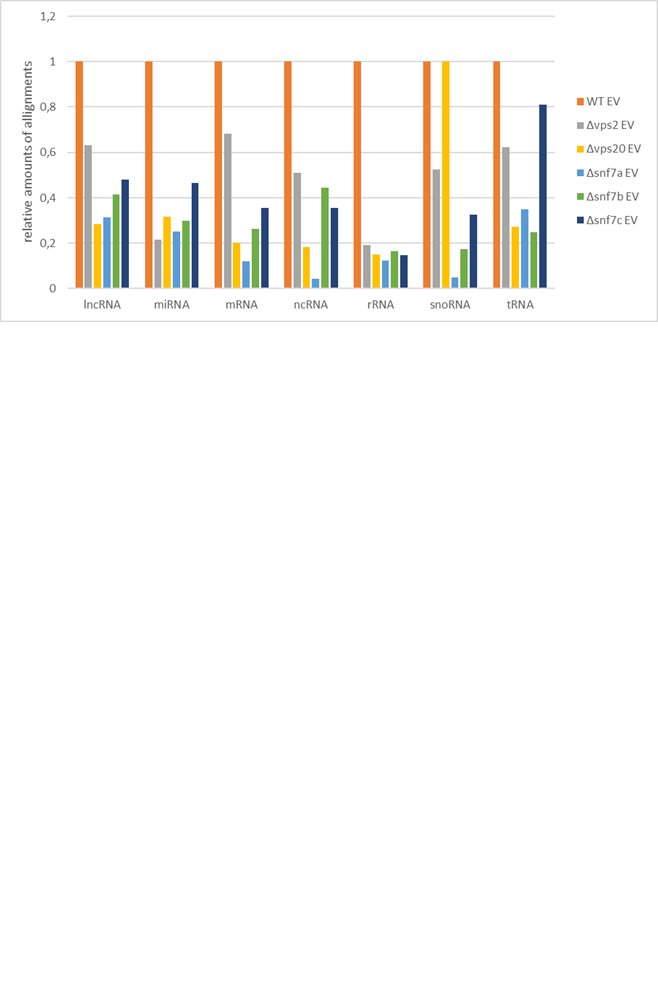

### Fig S5

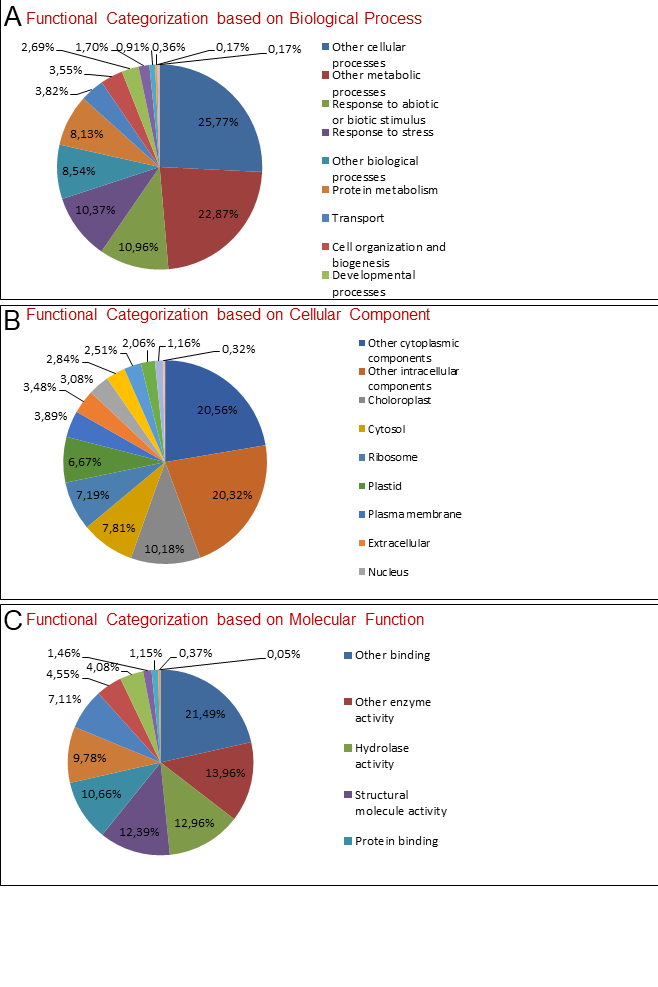
